# Rapidly evolving genes underlie *Aedes aegypti* mosquito reproductive resilience during drought

**DOI:** 10.1101/2022.03.01.482582

**Authors:** Krithika Venkataraman, Nadav Shai, Priyanka Lakhiani, Sarah Zylka, Jieqing Zhao, Margaret Herre, Joshua Zeng, Lauren A. Neal, Henrik Molina, Li Zhao, Leslie B. Vosshall

**Author notes:** Correspondence (KV), (LBV).

## Abstract

Female *Aedes aegypti* mosquitoes impose a severe global public health burden as vectors of multiple viral pathogens. Under optimal environmental conditions, *Aedes aegypti* females have access to human hosts that provide blood proteins for egg development, conspecific males that provide sperm for fertilization, and freshwater that serves as an egg-laying substrate suitable for offspring survival. As global temperatures rise, *Aedes aegypti* females are faced with climate challenges like intense droughts and intermittent precipitation, which create unpredictable, suboptimal conditions for egg-laying. Here we show that under drought-like conditions simulated in the laboratory, females retain mature eggs in their ovaries for extended periods, while maintaining the viability of these eggs until they can be laid in freshwater. Using transcriptomic and proteomic profiling of *Aedes aegypti* ovaries, we identify two previously uncharacterized genes named *tweedledee* and *tweedledum*, each encoding a small, secreted protein that both show ovary-enriched, temporally-restricted expression during egg retention. These genes are mosquito-specific, linked within a syntenic locus, and rapidly evolving under positive selection, raising the possibility that they serve an adaptive function. CRISPR-Cas9 deletion of both *tweedledee* and *tweedledum* demonstrates that they are specifically required for extended retention of viable eggs. These results highlight an elegant example of taxon-restricted genes at the heart of an important adaptation that equips *Aedes aegypti* females with “insurance” to flexibly extend their reproductive schedule without losing reproductive capacity, thus allowing this species to exploit unpredictable habitats in a changing world.

## INTRODUCTION

Extraordinary adaptations are ubiquitous across the animal kingdom within every habitat. Adaptations can be behavioral, physiological, or structural, and are evolutionarily selected to enable members of a species to persist by providing survival value. Ecosystems within which animals exist are in constant flux. When faced with changing habitats, animals must act flexibly and appropriately to survive and reproduce. For example, when the river-dwelling African lungfish (*Protopterus annectens*) experiences food and water scarcity during drought, it burrows into the dried riverbed, forming a cocoon with secreted mucus. It can survive for years while remaining metabolically dormant but within a week of rainfall, it reawakens and resumes normal metabolism (*1, 2*). Among birds, species like the blackcap (*Sylvia atricapilla*) exhibit genetically-encoded seasonal migratory behaviors that rapidly evolve in the face of changing resource availability, resulting in new migratory routes and destinations (*3–5*). Land mammals such as marsupials alter the timing of their reproductive cycle in response to offspring-derived cues (*6*). Tammar wallaby (*Macropus eugenii)* mothers can newly conceive while carrying previously birthed offspring in their pouches, but they developmentally arrest conceived embryos at the 100-cell stage, only resuming embryonic development after the pouch offspring has finished suckling and left to live independently (*6, 7*).

Mosquitoes that lay eggs at the edge of freshwater and go through an aquatic life cycle as larvae and pupae are very susceptible to fluctuating precipitation patterns and climate change-driven catastrophes like drought (*8–11*). Despite climate variations, *Aedes aegypti* mosquitoes are highly invasive on almost every continent and pose a serious, immediate, and growing threat to global public health (*12–15*). While biting multiple humans to obtain the protein-rich blood they require to develop each batch of eggs, these mosquitoes have evolved as efficient vectors of arboviral infections such as yellow fever, Zika, dengue, and chikungunya, and of parasitic infections such as lymphatic filariasis (*16*). Domestic strains of *Aedes aegypti* prefer hunting and biting humans over other vertebrate hosts (*17–19*), and prefer laying eggs on moist surfaces proximal to freshwater in natural and manmade containers found around human settlements (*20, 21*). Female *Aedes aegypti* typically mate once in their lifetime (*22*), storing sperm in specialized organs called the spermathecae from which sperm are released to fertilize eggs post-ovulation, as eggs are in transit through the reproductive tract *en route* to being laid (*23, 24*). Once laid at the edge of freshwater, eggs darken and harden, and embryogenesis occurs within the eggshell (*25*). After this, if conditions are suboptimal for hatching (*26*), a developmental arrest state called diapause is triggered to prevent embryo desiccation for up to 3-6 months. Embryos then hatch when pools of freshwater become available again, and when aquatic larval and pupal development can be completed before eclosion to the terrestrial adult stage (*27*).

The innate behaviors of an adult female *Aedes aegypti* mosquito are centered on the appropriate selection of a reproductive strategy that balances tradeoffs between internal energetic resources and external environmental conditions. Female mating, host-seeking, and egg-laying behaviors are inextricably linked, each proceeding only when the necessary “checkpoints” have been cleared (*28*). For example, females will not lay most, if any, of their eggs before they have mated (*29*). Females will suppress their attraction to hosts while eggs are developing and only restore attraction once eggs are laid (*30–35*). Females will not lay eggs both until and unless they locate freshwater, retaining them in their ovaries as needed (*21, 36, 37*). This interconnectedness of innate behaviors ensures that reproductive steps proceed in the order required for offspring survival. Because precise temporal control of egg-laying without loss of viability is an adaptation that maximizes the reproductive resilience and the fitness of *Aedes aegypti* females, understanding its basis will illustrate how this species is able to invade otherwise inhospitable ecological niches. Despite the importance of this question, little is known about how females are able to retain viable eggs in their ovaries during periods of prolonged drought.

Here we show that under drought-like conditions simulated in the laboratory, *Aedes aegypti* females will robustly retain eggs in their ovaries until freshwater is located. Under normal conditions when freshwater is plentiful, females will lay eggs 3-4 days after a blood meal. We restricted access to freshwater for 4-12 days post-blood meal. A considerable proportion of eggs laid after extended retention to at least 12 days post-blood-meal were viable, hatching at high rates. We identified two previously uncharacterized, tightly linked genes – here named *tweedledee* and *tweedledum* for the curious pair of characters in Lewis Carroll’s 1871 book, “Through the Looking-Glass and What Alice Found There” – that are adult female-specific and ovary-enriched in their expression. The expression of these genes is dramatically upregulated in the ovaries only during the period in which females retain eggs, and the genes are spatially limited to cells that encapsulate mature eggs. Both genes are taxon-restricted, with no detectable orthology except in *Aedes albopictus*, a similarly invasive disease vector mosquito species that is ~70 million years diverged from *Aedes aegypti* (*38*). In *Culex quinquefasciatus* and several *Anopheles* mosquito species, we identify “*conceptualogs*,” which we define as genes with no sequence homology to *tweedledee* or *tweedledum*, but which bear other featural similarities such as synteny, gene structure, gene size, and the presence of signal peptides in the predicted proteins. *Aedes aegypti tweedledee* and *tweedledum*, as well as the *Anopheles gambiae conceptualog*, are rapidly evolving genes within their respective species, and show strong signatures of positive selection. Using loss-of-function mutagenesis that disrupts both genes, we show that *Aedes aegypti tweedledee* and *tweedledum* are specifically required for extended retention of viable eggs under suboptimal drought conditions. Without *tweedledee* or *tweedledum*, mated, blood-fed *Aedes aegypti* females lose their reproductive “insurance,” such that when egg retention is triggered by restricted freshwater access due to drought-like conditions, most of the eggs they have matured no longer generate viable offspring if laid. Our results suggest that *tweedledee* and *tweedledum* play a crucial role in maintaining the reproductive resilience of female *Aedes aegypti* mosquitoes faced with fluctuating precipitation cycles and unpredictable drought-like conditions. This work thus illustrates a globally relevant example of rapidly evolving, taxon-restricted genes enabling an adaptation that allows *Aedes aegypti* mosquitoes to reproduce and thrive in otherwise inaccessible and inhospitable ecological niches.

## RESULTS

### Female innate reproductive behaviors are interconnected

To ensure that the timing and sequence of steps in a female *Aedes aegypti* mosquito’s reproductive cycle is appropriate for maximal reproductive output, the innate behaviors enabling access to blood meal sources, sperm, and freshwater egg-laying substrates are interconnected (Figure 1A). Under controlled laboratory conditions, consistent with previous results (*30–35*), we showed that virgin and mated females were both attracted to human hosts prior to blood-feeding, and mated females showed significantly stronger levels of attraction (Figure 1B). After blood-feeding, mated females suppressed their drive to hunt during egg development (Figure 1C–E). Mated females continued to suppress their host-seeking drive when they were forced to retain mature eggs in their ovaries, as in the absence of an egg-laying substrate, for moderate (6 days post-blood-meal, Figure 1C) or extended (12 days post-blood-meal, Figure 1D) periods. Using individual egg-laying vials (Figure S1A), we found that ~80% of mated females 6 days post-blood-meal completed egg laying within 3 hours of gaining access to freshwater (Figure S1B). Females showed restored attraction to humans within 2 hours of completing egg laying (Figure S1C–D). This shows that attraction to humans is only fully restored upon completion of egg-laying, at which time the female is ready to initiate a second cycle of reproduction. The dynamics of mosquito attraction to humans is similar across multiple cycles of reproduction (Figure 1E), which suggests that attraction to humans – a *de facto* protein-feeding drive – is strongly dependent on reproductive physiology. Virgin females, in contrast to mated females, laid few eggs when provided access to freshwater 6 days post-blood-meal (Figure 1F). Females that have both mated and have matured eggs within 3-4 days post-blood-meal must locate freshwater to lay their eggs (Figure 1G). If freshwater is not available, mated females will refrain from depositing their entire clutch of eggs (Figure 1G) (*21*). Blood-feeding and mating are thus decoupled – either one can occur first – but egg-laying, the ultimate step in a female’s reproductive sequence, is tightly coupled to both mating and blood-feeding and requires both to occur.

**Figure 1.**
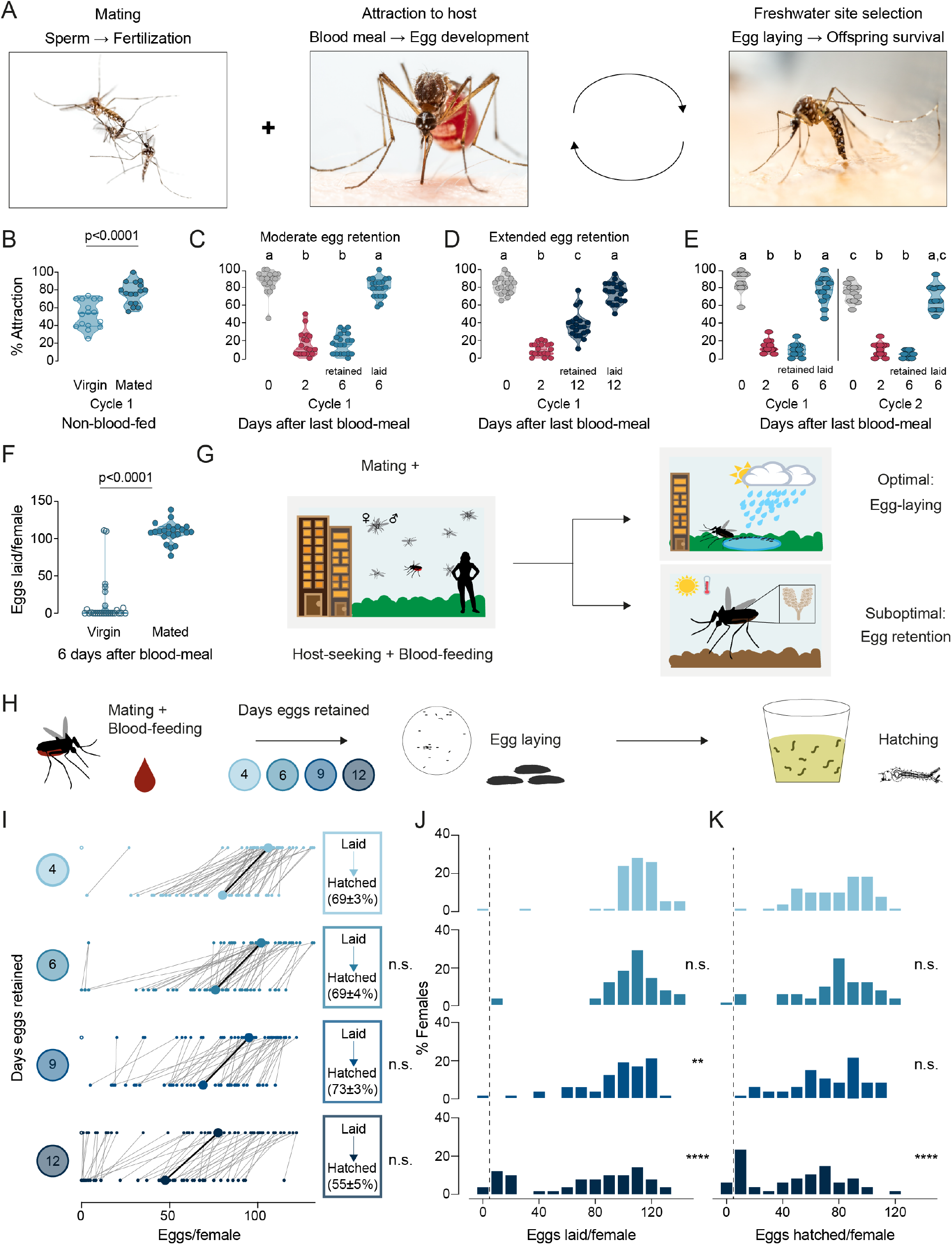
*Aedes aegypti* female reproduction is drought-resilient. (**A**) *Aedes aegypti* male and female mating (left), female blood-feeding from a human host (center), and female laying eggs in water (right). (**B-E**) Attraction of wild type females to a human forearm at the indicated reproductive state and cycle. Females are mated for all host-seeking experiments except where specified in (B). Each point represents a single trial with ~20 females, n=12-20 trials/group. Data are plotted as violin plots with median and 1st/3rd quartiles and showing all data points. Data labeled with different letters are significantly different: (B) Unpaired t-test, p<0.0001. (C) Kruskal-Wallis, Dunn’s multiple comparisons test, p<0.05. (D, E) one-way ANOVA, Tukey’s multiple comparisons test, p<0.05. (**F**) Number of eggs laid by mated or virgin females. Data represent eggs laid by a single female, n=22-26 females/group, shown as a violin plot with median and 1st/3rd quartiles with all data points. Mann-Whitney test, p<0.0001. (**G**) Schematic of a female mosquito’s reproductive decision point after egg maturation under optimal and suboptimal egg-laying conditions of freshwater abundance and scarcity, respectively. (**H**) Schematic of experiment to test effect of egg retention on egg laying and hatching. (**I**) Number of eggs laid by (top) and hatched from (bottom) single females that experienced the indicated egg retention periods. Females laying no eggs are depicted by open circles. Lines connect eggs laid by and hatched from the same individual. Larger circles and bold lines represent medians. Boxes show hatching rate (mean±S.E.M) from each egg retention group. n=46-50 females/group. (**J**, **K**) Distribution of eggs laid (J) and eggs hatched (K) after the indicated length of egg retention, analyzed from data in (I). Zero values are binned separately for each group. All other bins are groups of 10 starting with [1-10] and with closed/inclusive intervals. The groups for number of eggs laid (J), number of eggs hatched (K), and % hatched (I, boxes), respectively, at 6-, 9-, and 12-days post-blood-meal are compared to 4 days post-blood-meal to determine significant difference (Kruskal-Wallis, Dunn’s multiple comparison test; n.s.: not significant, p>0.05; **p<0.01; ****p<0.0001). Mosquito photographs (A): Alex Wild.

### Drought induces extended retention of viable eggs

When a mated female mosquito has converted blood meal nutrients into mature eggs over 3-4 days, she must not only make the decision of where to lay her eggs, but she must also appropriately time her egg-laying decisions to ensure maximal offspring survival. We measured egg retention in the laboratory by simulating drought-like conditions of varying durations (Figure 1H). Females that engorged on a full blood meal laid ~100-110 eggs at the edge of freshwater 4 days after a blood meal, of which ~70% hatched (Figure 1I–K). Even though the number of eggs laid decreased with increasing length of egg retention, the proportion of viable eggs remained consistently high even after extended egg retention to at least 12 days post-blood-meal (Figure 1I–K). These results show that wild type *Aedes aegypti* mosquitoes demonstrate remarkable reproductive resilience during drought by retaining viable eggs until freshwater becomes available.

### Candidate regulators of viable egg retention are ovary-enriched with temporally restricted expression

How female *Aedes aegypti* mosquitoes carrying mature eggs in their ovaries maintain the potential for subsequent fertilization, laying, and hatching of their eggs after different lengths of retention remains unexplored. To identify candidate genes regulating the retention of viable eggs in *Aedes aegypti* ovaries post-maturation, we used bulk RNA-sequencing (RNA-seq) to profile ovaries across 11 different time-points in their first cycle of reproduction (Figure 2A). Principal component analysis (PCA) of the ovary RNA-seq dataset shows that replicates within each reproductive stage cluster together, and that principal components 1 and 2 (PC1 and PC2) separate the reproductive stages from each other (Figure 2B). These data reflect the major transcriptional changes in the ovary across the reproductive cycle and highlight that each stage is distinctly and tightly regulated.

**Figure 2.**
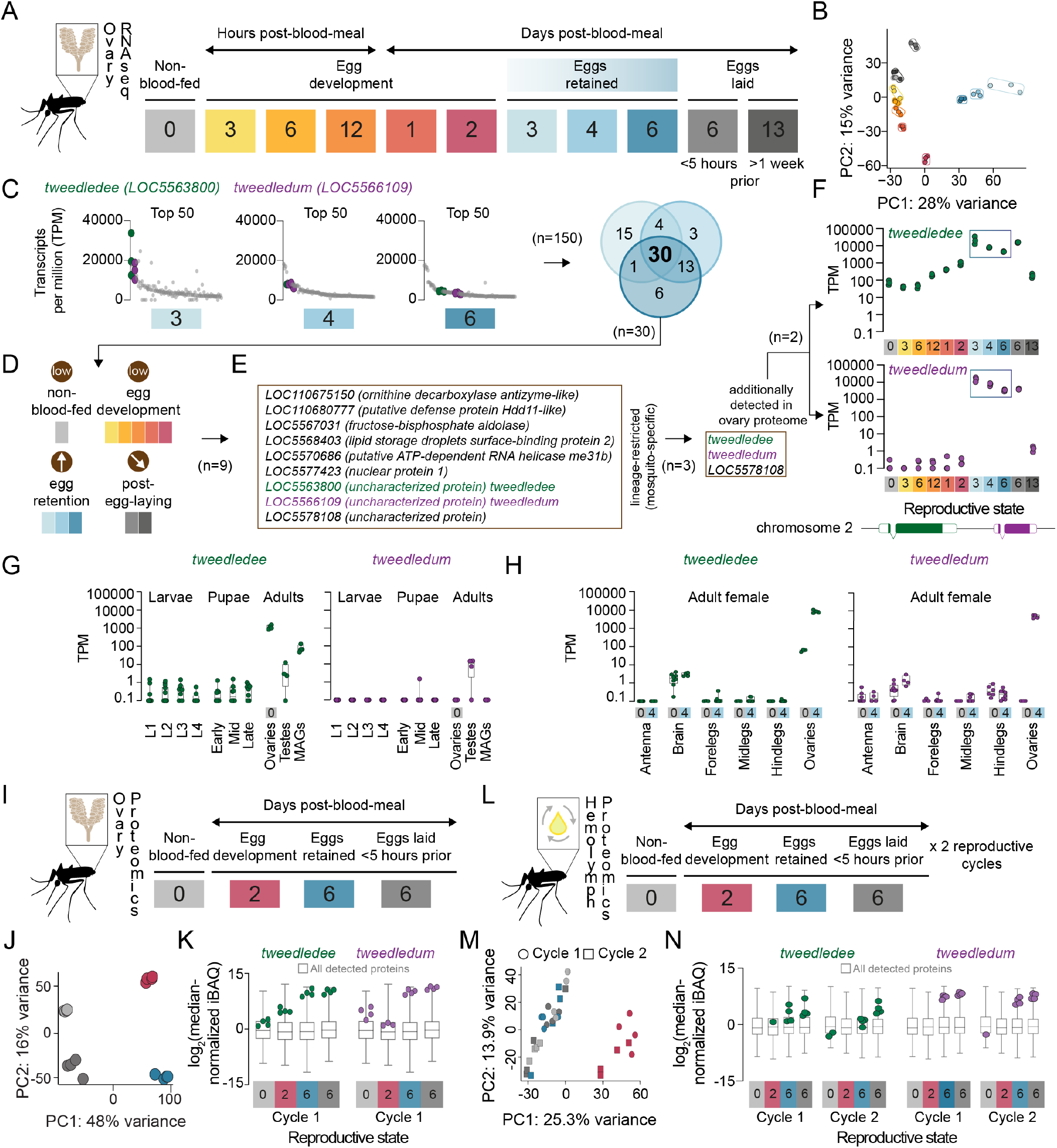
*tweedledee* and *tweedledum* are ovary-enriched and upregulated during egg retention. (**A**) Reproductive time-points sampled for bulk ovary RNA-seq, n=3 replicates/group, 11 groups. (**B**) Principal component analysis (PCA) of DESeq2-normalized, transformed counts from ovary RNA-seq. (**C**) Top 50 most abundant transcripts ranked by median transcripts per million (TPM) for egg retention groups, 3-, 4-, and 6-days post-blood-meal (total=150 transcripts). Gray dots represent replicates for each transcript in the top 50, green dots indicate *tweedledee*, and purple dots represent *tweedledum.* Venn diagram of the number of overlapping transcripts between the top 50 genes across egg retention groups, with 30 transcripts shared between all three groups. (**D**) Logic of expression pattern for selecting candidate genes expected to underlie robust retention of viable eggs. (**E**) Genes with expression dynamics that follow filtering criteria in D. (**F**) Transcript expression pattern in the ovaries of *tweedledee* (top) and *tweedledum* (bottom), which were filtered as final candidates based on their additional detection in the ovary proteome (see Figure 2K) and their being taxon-restricted. The blue rectangle indicates the period of egg retention. (**G**) TPM values for *tweedledee* (left) and *tweedledum* (right) during larval, pupal, and adult stages of development [data from (*40*)], MAGs = male accessory glands, n=4-13 replicates/group. (**H**) TPM values for *tweedledee* (left) and *tweedledum* (right) in adult female tissues (data originally from (*41*), reanalyzed in (*40*), n=3-8 replicates/group. (**I**) Reproductive time-points sampled for ovary proteomics, n=4 replicates/group, 4 groups. (**J**) PCA of iBAQ values from ovary proteomics. (**K**) Distribution of iBAQ values as a metric of abundance for all proteins detected per group in ovary proteomics. Overlaid green dots represent individual replicate values for *tweedledee* and purple dots represent replicates for *tweedledum.* All values are pre-imputation and represent log^2^-transformed median iBAQ signals normalized by subtracting the median iBAQ signal for the group. (**L**)Reproductive time-points sampled for hemolymph proteomics, n=4 replicates/group, 8 groups. (**M**) PCA of iBAQ values from hemolymph proteomics. (**N**) Distribution of iBAQ values as a metric of abundance for all proteins detected per group in hemolymph proteomics. Overlaid green dots represent individual replicate values for *tweedledee* and purple dots represent replicates for *tweedledum.* All values are pre-imputation and represent log^2^-transformed median iBAQ signals normalized by subtracting the median iBAQ signal for the group. Box plots in G, H, K, N: median, 1^st^/3^rd^ quartile, minimum to maximum whiskers.

Ovaries carrying mature eggs occupy much of the female mosquito’s abdomen, requiring redirection of her energy resources towards maintaining her eggs. Therefore, we hypothesized that candidate regulators of viable egg retention would be abundantly expressed across egg retention time-points (Figure 2C), with specific upregulation at time-points when eggs are retained compared to pre-blood meal or during egg development (Figure 2D). We expected that post-egg-laying, the expression of these genes would eventually decline as the female transitions out of her reproductive state (Figure 2D). We speculated that, given the distinct natural histories and diverse egg-laying strategies across insects (*20, 37, 39*), genes regulating viable egg retention in *Aedes aegypti* may be taxon-restricted within mosquitoes (Figure 2E).

Filtering based on these steps, and by confirming the presence of candidates in an ovary proteome dataset that we generated (see Figure 2K), we identified two genes that satisfy our criteria as candidate regulators of viable egg retention. These previously uncharacterized genes, *LOC5563800* and *LOC5566109*, which we named *tweedledee* and *tweedledum* respectively, show similar and striking patterns of regulation in our transcriptomic dataset (Figure 2F). *tweedledee* is expressed in females before a blood meal and during egg development, but its expression increases 3 orders of magnitude during periods of egg retention. The regulation of *tweedledum* is even more remarkable. It is present at less than 1 transcript per million (TPM) at non-blood-fed and egg production stages but rises to 10,000 TPM during egg retention.

Using published transcriptomes of developmental stages (*40*) and adult tissues (*41*) from *Aedes aegypti* mosquitoes, we confirmed that *tweedledee* and *tweedledum* are adult-specific and female-enriched (Figure 2G). They both show ovary-enriched expression with strong upregulation post-egg maturation in the females (Figure 2H), with some expression in male reproductive tissues (Figure 2G). In addition to specific upregulation during egg retention, *tweedledee* shows basal, constitutive expression across several conditions and tissues, whereas the spatiotemporal expression of *tweedledum* is more tightly restricted (Figure 2F–H). These data in adult females show exquisite specificity of *tweedledee* and *tweedledum* expression in ovaries bearing mature eggs, strengthening the possibility that the genes are candidate regulators of viable egg retention.

We next performed quantitative proteomics profiling of the female ovaries across a subset of reproductive time-points corresponding to non-blood-fed, egg development, egg retention, and post-egg-laying states (Figure 2I). PCA shows that replicates within each stage again clustered together as in the RNA-seq dataset, and all reproductive states formed distinct clusters in PC1 and PC2, reflective of the ovaries being tightly and distinctly controlled across these different reproductive states (Figure 2J). Both tweedledee and tweedledum protein were notably upregulated at the egg retention phase of the reproductive cycle (Figure 2K). Because tweedledee and tweedledum expression levels remain high in the ovaries when sampled <5 hours post-egg-laying when all mature eggs have been laid, we speculate that these genes are expressed in somatic tissues in the ovary (Figure 2K).

*tweedledee* and *tweedledum* are predicted to encode proteins with N-terminal signal peptides. To test if they are secreted, we profiled the proteome of the circulating hemolymph, the insect equivalent of blood. We collected hemolymph samples across non-blood-fed, egg development, egg retention, and post-egg-laying states in the first and second cycles of reproduction (Figure 2L). The hemolymph is in close apposition to the ovaries, and its contents during distinct reproductive time-points reflect interorgan communication (*42–44*). PCA of the hemolymph proteome showed that at 2 days post-blood-meal, the composition of the circulating fluid is most distinct (separated by PC1) from other profiled time-points (Figure 2M). These findings are consistent with our expectations, as this is the only time-point profiled during which eggs are likely to still be maturing and during which the hemolymph is therefore transporting components for egg maturation (*45, 46*). Notable examples of hemolymph-transported proteins include the vitellogenins (yolk protein precursors), which we detect in our dataset (https://doi.org/10.5281/zenodo.5945525). These proteins are synthesized in the fat body, an analog of the vertebrate liver, and transported via the hemolymph to the ovaries where they are packaged into maturing eggs (*45, 47*). We detected tweedledee and tweedledum protein in the hemolymph and found that they were both strongly upregulated in each of the reproductive cycles during egg retention and within 5 hours of egg-laying compared to pre-blood-meal, during egg development, or >1-week post-egg-laying (Figure 2N). These data together suggest that somatic ovary cells secrete tweedledee and tweedledum, and that their expression and secretion into the circulating hemolymph are both tightly regulated.

### *tweedledee* and *tweedledum* are expressed in cells encapsulating mature eggs

Within the ovaries, mature eggs are housed in individual chambers/follicles, encapsulated within a membrane of follicular epithelial cells (*48, 49*). At the point of egg-laying, mature eggs transit out of their individual chambers and enter the calyx, a continuous tube through the center of the ovary connected to the oviducts (*49, 50*). Eggs transit into the oviducts and are fertilized in the reproductive tract by sperm released from sperm storage organs, the spermathecae, before being ejected through the ovipositor (*24, 49, 50*).

Using whole-mount ovary fluorescence RNA *in situ* hybridization we show that *tweedledee*, but not *tweedledum* transcripts are detectable in non-blood-fed ovaries (Figure 3A). *tweedledee* expression in non-blood-fed ovaries is restricted to calyx cells, and it is markedly absent in the primary follicles (Figure 3A). The primary follicles are comprised of seven nurse cells and an oocyte surrounded by somatic follicular epithelial cells (*51*). Once the female consumes a blood meal, the primary follicle develops into an egg, with the surrounding follicular epithelial cells secreting eggshell proteins and other components onto it (*51, 52*). The oocyte is characteristically marked by *vitellogenin receptor* (*LOC5569465*) expression (Figure 3A,B,D,F). The *vitellogenin receptor* gene enables receptor-mediated endocytosis of yolk precursor proteins into the egg after a blood meal (*53*).

**Figure 3.**
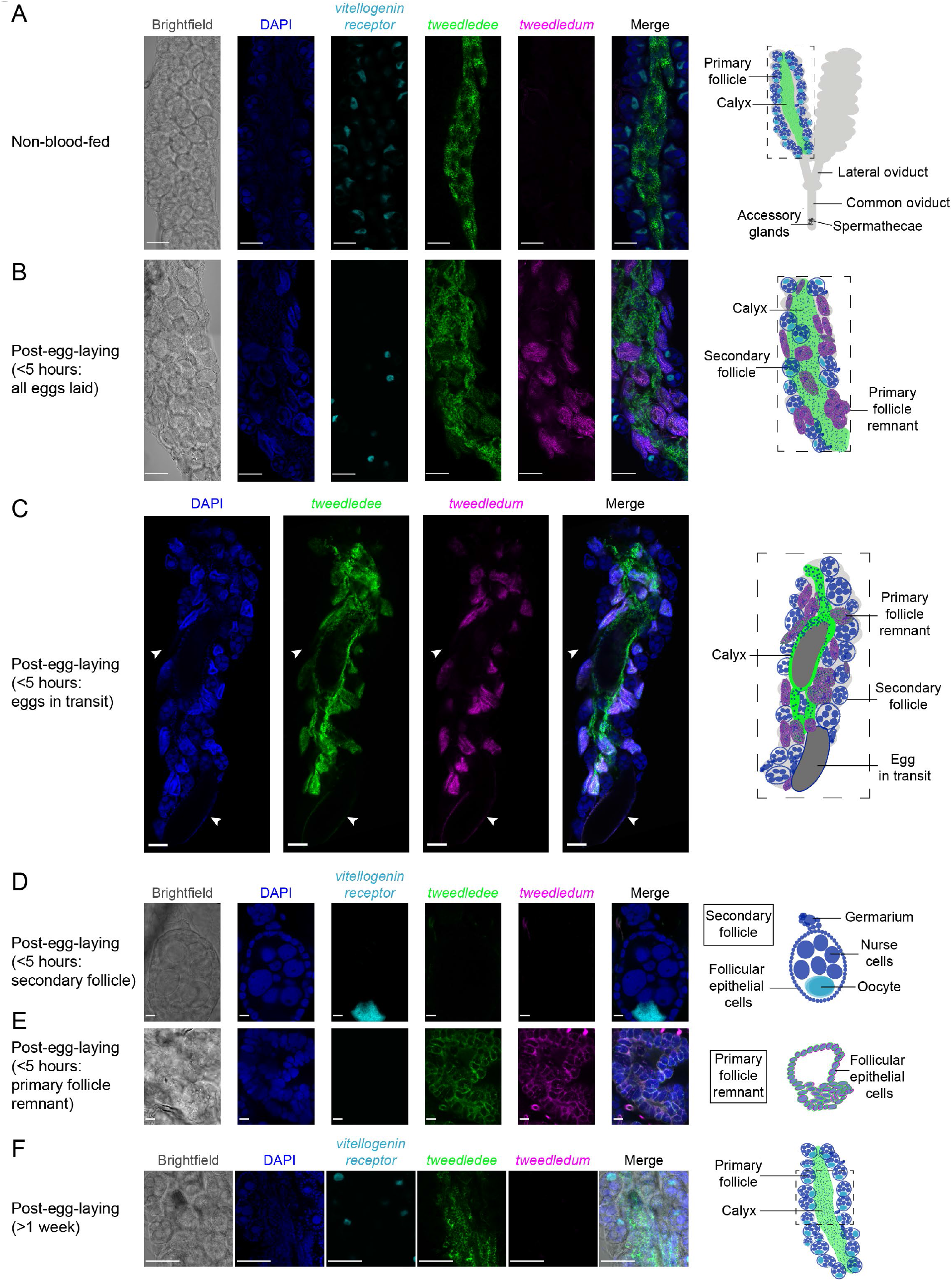
*tweedledee* and *tweedledum* are expressed in cells encapsulating mature eggs. (**A**) Left: Single confocal section of whole-mount fluorescence RNA *in situ* hybridization of a non-blood-fed ovary with the indicated probes. Right: cartoon of a pair of ovaries and the female reproductive system, with left ovary representing a cross-section of one ovary. (**B, C**) Left: Single confocal section of whole-mount fluorescence RNA *in situ* hybridization of an ovary <5 hours post-egg-laying with all eggs laid (B) or with two eggs in transit indicated by white arrows (C) with the indicated probes. Right: cartoons representing cross-section through a post-egg-laying ovary with probe expression patterns depicted in different ovary structures. (**D, E**) Left: Single confocal section of whole-mount fluorescence RNA *in situ* hybridization of an ovary <5 hours post-egg-laying with the indicated probes showing a secondary follicle ready to develop into an egg upon consumption of a second blood meal (D) and a post-egg-laying follicle that is the remnant of a primary follicle which previously contained an egg (E). Right: cartoons depicting *tweedledee* and *tweedledum* expression pattern uniquely in the post-egg-laying follicle/primary follicle remnant (E), but not in the secondary follicle expressing vitellogenin receptor (D). (**F**) Left: Single confocal section of whole-mount fluorescence RNA *in situ* hybridization of an ovary >1-week post-egg-laying with the indicated probes. Right: cartoon of a single ovary cross-section. Scale bars: 100 μm in A-C, F and 10 μm in D-E.

Because of technical limitations of performing RNA *in situ* hybridization on intact ovaries both during egg development and during retention of fully mature eggs due to optical opacity of the ovary and difficulties with probe penetration, we utilized ovaries 6 days post-blood-meal within 5 hours of egg-laying to identify the cells expressing *tweedledee* and *tweedledum* (Figure 3B-E). Since ovary RNA-seq data suggest both transcripts are abundantly expressed <5 hours post-egg-laying (Figure 2F), we postulated that this time-point would allow us to identify which cells express *tweedledee* and *tweedledum*. Ovaries collected within 5 hours of egg-laying have two different types of egg follicles (Figure 3B–E): the remnants of primary follicles which held mature eggs prior to laying and a secondary follicle that was previously attached to the primary follicle, and that is ready to develop into a new egg upon consumption of a second blood meal (*49, 51, 54*). *tweedledee* was detected in the calyx through which eggs transit (Figure 3C) as well as in the follicular epithelial cells of the primary follicle remnants (Figure 3B–E). *tweedledum* was also expressed in the follicular epithelial cells of primary follicle remnants, and solely in these cells together with *tweedledee* (Figure 3B–E). Notably, neither of the transcripts were expressed in secondary follicles (Figure 3B–E).

We additionally examined *tweedledee* and *tweedledum* expression >1-week post-egg-laying when the gross morphology of the ovary resets and bears closer resemblance overall to non-blood-fed ovaries. At this time-point, *tweedledum* expression was once again undetectable, and *tweedledee* was exclusively expressed in the calyx (Figure 3F). The patterns of *tweedledee* and *tweedledum* expression detected using RNA *in situ* hybridization in non-blood-fed ovaries (Figure 3A) and in ovaries <5 hours (Figure 3B–E) or >1-week post-egg-laying (Figure 3F) validate the expression patterns from the respective time-points in the ovary RNA-seq data (Figure 2F). Overall, the robust expression of *tweedledee* and *tweedledum* in the follicular epithelial cells of primary follicle remnants and the added expression of *tweedledee* in the calyx suggests that these genes, either independently or together, are poised to play a role in protecting eggs specifically during egg retention and while they are *en route* to being laid.

### *tweedledee* and *tweedledum* are linked, taxon-restricted, and rapidly evolving

We next to turned to the *Aedes aegypti* genome for clues on the function and evolutionary origin of *tweedledee* and *tweedledum*. The genes are located next to each other on chromosome 2, and both have a short first exon and a longer second exon (Figure 4A). These genes are predicted to encode small proteins (tweedledee: 216 amino acids; tweedledum: 116 amino acids), both with N-terminal signal peptides but no other known domains (Figure 4B). Although similar in many respects, the two genes and their encoded proteins bear no sequence similarity to each other. We calculated the guanine+cytosine (GC) content for all protein-coding genes in the *Aedes aegypti* genome. This metric is indicative of gene and transcript thermal stability, and thus also an important determinant shaping interactions between a species and its environment (*55*). Compared to all protein-coding genes, *tweedledee* (50% GC) and *tweedledum* (48% GC) fall in the 94^th^ and 89^th^ percentile, respectively (Figure 4C). Within the distribution of protein-coding genes containing a predicted signal peptide, the percentiles for *tweedledee* and *tweedledum* remain similar at 95^th^ and 88^th^, respectively (Figure 4C). We next calculated the proportion of each amino acid residue in tweedledee and tweedledum and compared it to the average proportion of each amino acid residue across all proteins in the *Aedes aegypti* genome that contain a predicted signal peptide (Figure 4D). In all cases, we performed comparisons on proteins in their functional secreted form, with signal peptides cleaved *in silico*. Both tweedledee and tweedledum share compositional biases with each other relative to other secreted proteins encoded by the *Aedes aegypti* genome. They both show an underrepresentation of leucine, threonine, and glycine, an overrepresentation of aspartate, glutamate, alanine, valine, and serine, and entirely lack cysteine, tyrosine, and tryptophan (Figure 4D).

**Figure 4.**
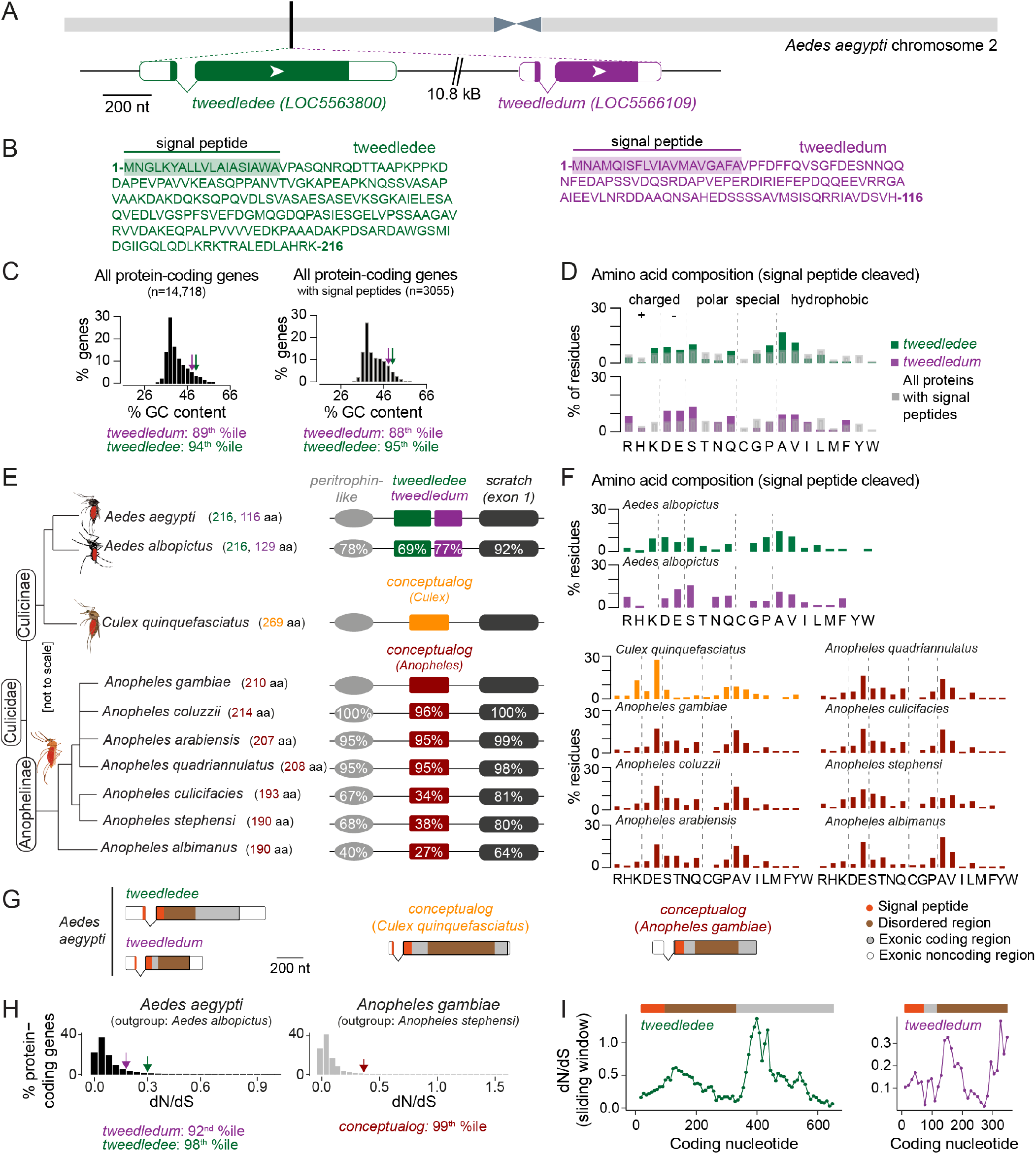
*tweedledee* and *tweedledum* are linked, taxon-restricted, syntenic, and rapidly evolving genes. (**A**) Chromosomal location and gene structure of *tweedledee* and *tweedledum*. (**B**) Amino acid sequences of tweedledee (left) and tweedledum (right) with predicted N-terminal signal peptides indicated. (**C**) GC content of all protein-coding genes in the *Aedes aegypti* (AaegL5) genome (left) and of all protein-coding genes with predicted signal peptides (right), with *tweedledee* and *tweedledum* indicated by arrows. (**D**) Amino acid composition of *Aedes aegypti* tweedledee and tweedledum, as compared to mean percent residue for 3,040 proteins with predicted signal peptides in the *Aedes aegypti* genome, calculated after signal peptide cleavage. Syntenic loci in *Aedes*, *Culex*, and *Anopheles* mosquito species are shown (not to scale), ordered by the topology of the mosquito phylogenetic tree. The protein length of tweedledee, tweedledum, or the conceptualog is shown in parentheses next to the species name. Protein sequence identity is shown for each gene as calculated using a reference species for each genus, either *Aedes aegypti* or *Anopheles gambiae*. For *scratch*, protein sequence identity was calculated by aligning exon 1 of each species due to a fragmented annotation in multiple reference genomes (see Methods). Accession numbers for all genes are at https://doi.org/10.5281/zenodo.5945525. Amino acid composition of tweedledee *and* tweedledum in *Aedes albopictus* and of the conceptualogs in *Culex* and *Anopheles* species. Gene structures of *Aedes aegypti tweedledee* and *tweedledum* and the *conceptualog* in *Culex quinquefasciatus* and *Anopheles gambiae* are shown to scale with signal peptide and disordered domains annotated. The 3’UTR of *Anopheles gambiae conceptualog* is lacking in the current genome annotation. (**H**) The distribution of dN/dS values for 8,030 protein-coding genes in *Aedes aegypti* and 9,958 protein-coding genes in *Anopheles gambiae*. *tweedledee*, *tweedledum* and the *conceptualog* are shown with arrows. (**I**) dN/dS values were calculated for a 102-nucleotide sliding window size of 34 nucleotides each with a 3 amino acid overlap across the coding sequence of *Aedes aegypti tweedledee* and *tweedledum*. Coding sequences were aligned to orthologs in *Aedes albopictus*.

To explore the evolutionary history and origin of these genes, we searched for putative homologs. Using BLASTp, the only orthologs identifiable in Genbank for both tweedledee and tweedledum with E-values < 0.05 are in *Aedes albopictus*, another invasive mosquito vector ~70 million years diverged from *Aedes aegypti* (*38*). In both *Aedes aegypti* and *Aedes albopictus, tweedledee* and *tweedledum* or their respective orthologs are flanked by two conserved genes, *peritrophin-like* and *scratch* (annotated as *escargot* in *Aedes aegypti*, see Methods) (Figure 4E). Using *peritrophin-like* and *scratch* as “anchor” genes, we searched for other syntenic loci potentially containing *tweedledee* or *tweedledum* homologs in other mosquito species (Figure 4E). We found syntenic loci in several other mosquitoes, but not in any non-mosquito species, including *Drosophila melanogaster* flies. The *Drosophila melanogaster scratch* gene is located on chromosome 3L and is not in close proximity to any *peritrophin-like* genes. There are several genes adjacent to *Drosophila melanogaster scratch*, but none have the gene or protein structure of *tweedledee* and *tweedledum*. In *Culex quinquefasciatus, Anopheles gambiae*, and several other *Anopheles* mosquito species both within and outside of the *Anopheles gambiae* complex, there are syntenic loci with conserved *peritrophin-like* and *scratch* genes (Figure 4E). In these *Culex* and *Anopheles* cases, the conserved genes flank a single uncharacterized gene (Figure 4E). We termed these single uncharacterized genes in non-*Aedes* mosquitoes “*conceptualogs*,” since they bear no sequence homology to *tweedledee* or *tweedledum* in *Aedes*, but they are like *tweedledee* and *tweedledum* in many other aspects. First, they are all two exons long, with a short first exon and a longer second exon. Second, they are predicted to encode proteins of similar length ranging between 190 and 269 amino acids, and third, they are predicted to contain N-terminal signal peptides. Ordered by the topology of the mosquito phylogenetic tree, the protein sequences of tweedledee and tweedledum in *Aedes* or of the conceptualogs in *Anopheles* diverge more rapidly than the protein sequences of their flanking anchor genes within their respective genera (Figure 4E). In comparing the amino acid content of all conceptualogs (with signal peptides cleaved) to each other and to the *Aedes* tweedledee and tweedledum, we observed several similarities despite the rapid protein sequence divergence: all genes have no or very few cysteine or tryptophan residues, and an overrepresentation of glutamate and alanine (Figure 4F). Exonic sequences of *tweedledee*, *tweedledum*, and the *conceptualogs* in *Culex* and *Anopheles* also show strong similarities in the relative locations of their signal peptides and disordered domains as predicted by SignalP and IUPred2A, respectively (Figure 4G).

To assess whether the molecular evolution of *Aedes aegypti tweedledee* and *tweedledum* relative to the outgroup, *Aedes albopictus*, and of the *Anopheles gambiae conceptualog* relative to the outgroup, *Anopheles stephensi*, is adaptive, we computed the ratio of non-synonymous (dN, amino acid-altering) to synonymous (dS, silent) mutations at each site (*56*). By calculating the distribution of dN/dS values for all protein-coding genes with unique outgroup orthologs in the *Aedes aegypti* and *Anopheles gambiae* genomes, we found that *tweedledee*, *tweedledum*, and the *conceptualog* are in the 98^th,^ 92^nd^, and 99^th^ percentile, respectively (Figure 4H). This suggests that compared to most protein-coding genes in mosquitoes, amino acid-altering mutations are more likely to reach fixation for *tweedledee*, *tweedledum*, and the *Anopheles conceptualog*. A sliding-window analysis of dN/dS values across the coding sequences of *Aedes aegypti tweedledee* and *tweedledum* revealed that these high gene-wide dN/dS values are likely driven by rapid sequence divergence in specific regions around the middle of the gene (Figure 4I). These analyses together suggest that *tweedledee, tweedledum*, and the *conceptualogs* likely shared a common ancestor, and that the genes are evolving rapidly under strong selective pressure across the mosquito phylogeny.

### *tweedledee* and *tweedledum* are required for retention of viable eggs during drought

Under fluctuating climate conditions of intermittent precipitation, retaining viable eggs for extended durations may be an adaptive reproductive strategy for *Aedes aegypti* females. To test whether *tweedledee* and *tweedledum* are required under drought-like conditions for females to retain viable eggs for extended periods after blood feeding, we used CRISPR-Cas9 to generate a large deletion at the *tweedledee* and *tweedledum* locus, here referred to as *Δdeedum* double mutants (Figure 5A, Supplementary Figure S2A). The 11.7 kb deletion starts within *tweedledee* exon 2 and ends in exon 2 of *tweedledum* (Figure 5A). The gene fusion resulting from the large deletion and several additional indels is predicted, *in silico*, to encode a protein with amino acids 1-53 of *tweedledee* conserved before the breakpoint junction, following which a frameshift is introduced that leads to fusion with 68 missense amino acids before a stop codon (Figure 5A). This deletion event in *Δdeedum* is predicted to produce null mutations in both *tweedledee* and *tweedledum*.

**Figure 5.**
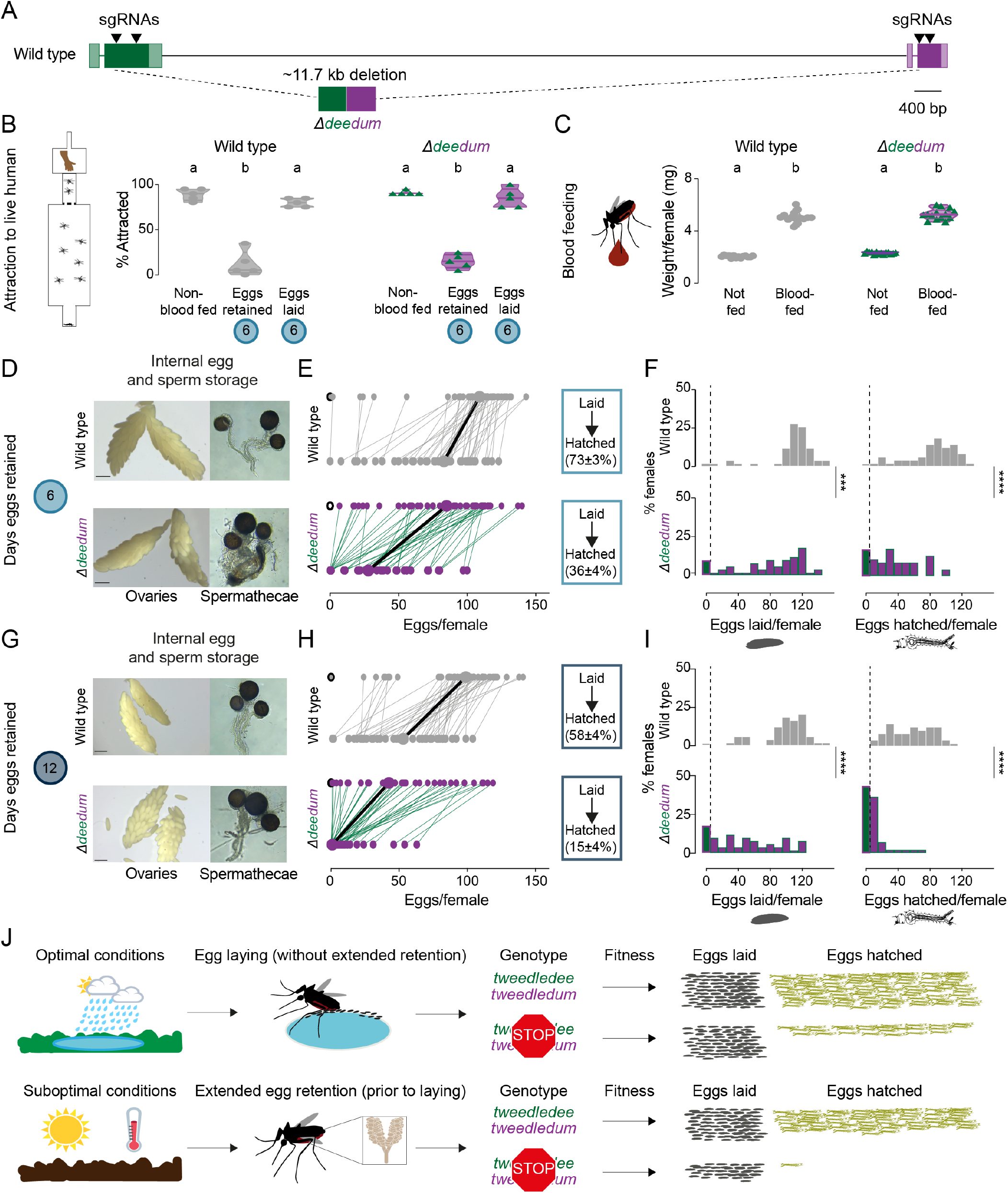
*tweedledee* and *tweedledum* are required for reproductive resilience during drought. (**A**) Schematic of *Δdeedum* mutant that deletes both *tweedledee* and *tweedledum*. (**B**) Attraction of wild type and *Δdeedum* mutant females to a human forearm. Data are plotted as violin plots with median and 1st/3rd quartiles and showing all data points. Each point represents a single trial with ~20 females, n=5 trials/group. Significantly different groups are indicated by different letters (one-way ANOVA, Holm-Šídák’s multiple comparisons test, p<0.0001). (**C**) Averaged weights of 5 females of the indicated genotype not fed or blood fed, n=14 groups of 5 females per group. Data are plotted as violin plots with median and 1st/3rd quartiles and showing all data points. Significantly different groups are indicated by different letters (one-way ANOVA, Tukey’s multiple comparisons test, p<0.0001). (**D, G**) Photographs of ovaries (left) and spermathecae with filled sperm (right) from wild type and *Δdeedum* females 6 days (D) or 12 days (G) post blood-meal with eggs retained. (**E, H**) Number of eggs laid by (top) and hatched from (bottom) single wild type and *Δdeedum* mutant females 6 days (E) and 12 days (H) post blood-meal, depicting moderate and extended egg retention, respectively. Females laying no eggs are depicted by open circles. Lines connect eggs laid by and hatched from the same individual. Larger circles and bold lines represent medians. Boxes show (mean S.E.M) of % eggs hatched from each egg retention group, n = 48-50 females/ group. (**F, I**) Distribution of eggs laid (left) and eggs hatched (right) after egg retention in wild type and *Δdeedum* mutant females 6 days (F) or 12 days (I) post-blood-meal. Zero values are binned separately for each group. All other bins are groups of 10 starting with [1-10] and with closed/inclusive intervals. (F, I) The groups between each genotype for eggs laid and eggs hatched respectively were compared at each of the time points to determine significant difference (Mann-Whitney tests, *** p<0.001; **** p<0.0001). Distributions in (F) are analyzed from data in (E) and distributions in (I) are analyzed from data in (H). (**J**) Summary of *tweedledee* and *tweedledum* function in drought resilience of female *Aedes aegypti* mosquitoes.

To characterize the reproductive behaviors of *Δdeedum* double mutant females compared to wild type females, we mated them to sibling males of their respective genotypes. To assess the general health of females, we tested their level of attraction to human hosts (Figure 5B) and their blood meal consumption (Figure 5C). Like wild type females, *Δdeedum* double mutant females were strongly attracted to a live human arm in a single stimulus olfactometer assay (Figure 5B). They approximately doubled their body weight from engorging on a blood meal (Figure 5C), and when presented with the same live arm stimulus 6 days following the blood meal while retaining eggs, they showed a suppressed host-seeking drive like wild type females (Figure 5B). Both wild type and *Δdeedum* females restored attraction to human hosts by 6 days after the blood meal if they had been provided freshwater to lay eggs 3-5 days after the blood meal (Figure 5B). Together, these host-seeking and blood-feeding results suggest that *Δdeedum* mutants are healthy, and that loss of *tweedledee* and *tweedledum* together does not affect attraction to human hosts, modulation of attraction following a blood meal and following egg-laying, or the ability to engorge on a full blood meal – all crucial behavioral checkpoints for reproductive success.

We next asked whether *Δdeedum* double mutant females have morphologically healthy ovaries and spermathecae with visually normal eggs and sperm, respectively. *Δdeedum* females that consumed a full blood meal developed mature eggs and retained them for at least 12 days after the blood meal in their ovaries. There were no grossly observable morphological defects in *Δdeedum* eggs or ovaries compared to wild type when ovaries were dissected and photographed 6 days (Figure 5D) or 12 days (Figure 5G) after the blood meal. Spermathecae, the organs specialized for sperm storage following a single mating event contained sperm that appeared motile in both wild type and *Δdeedum* mutants at both time-points (Figure 5D,G).

We then tested the egg retention and egg-laying behaviors of *Δdeedum* females compared to wild type to assess how the mutants compare to wild type in their reproductive resilience during drought. We blood fed wild type and *Δdeedum* mutant females, and withheld access to a freshwater substrate for either 6 days (Figure 5E,F) or 12 days (Figure 5H,I), corresponding to moderate or extended drought-like conditions. When we provided freshwater 6 days after the blood meal, 98% of wild type females compared to 90% of *Δdeedum* females laid at least one egg (Figure 5E,F). While wild type females laid a median of 109 eggs, *Δdeedum* mutant females laid a median of 85 eggs. Of the females that laid any eggs, 98% of the wild type females compared to 82% of *Δdeedum* females produced at least one viable offspring (Figure 5E,F). Of the eggs laid by the respective genotypes, 73% of wild type eggs hatched, while only 36% of *Δdeedum* eggs hatched (Figure 5E,F). Deleting *tweedledee* and *tweedledum* therefore had a considerable effect on egg viability during moderate egg retention.

If instead we withheld freshwater for 12 days after the blood meal before providing it to females, 98% of wild type females still laid at least one egg compared to 82% of *Δdeedum* females (Figure 5H,I). Of the females that laid eggs, 100% of the wild type females still produced at least one viable offspring compared to only 56% of *Δdeedum* females (Figure 5H,I). Wild type animals laid a median of 98 eggs of which 58% hatched (Figure 5H,I). *Δdeedum* females laid a median of 43 eggs after this extended retention, but in stark contrast to wild type, only 15% of the eggs laid hatched (Figure 5H,I). Therefore, as the duration of drought increased, females lacking *tweedledee* and *tweedledum* very significantly lost their ability to remain reproductively resilient. At both 6 days (Supplementary Figure S2B) and 12 days (Supplementary Figure S2C) post-blood-meal, heterozygous females laid a similar number of eggs to wild type females, indicating that the *Δdeedum* phenotype is recessive.

*Aedes aegypti* females undergo profound changes in physiology and behavior upon mating (*29, 57–59*). To ask if the genotype of the male with which the female had mated had an influence on these female reproductive resilience phenotypes, we mated both wild type and *Δdeedum* females to wild type males instead of sibling males of their respective genotypes. We tested the number of eggs laid by females after extended retention and found that *Δdeedum* females laid significantly fewer eggs compared to wild type females (Supplementary Figure S2D). These data show that the decreased fitness after egg retention seen in *Δdeedum* females is a maternally-derived phenotype.

In this study we generated a deletion that disrupted both *tweedledee* and *tweedledum*. This left open the question of whether both genes contribute to reproductive resilience during drought. To date, we have been unable to establish a homozygous *Δdee* single mutant strain. However, we recovered a *Δdum* single mutant (Supplementary Figure S3A–B) and observed that it had a phenotype largely overlapping with that of *Δdeedum* (Supplementary Figure S3C–I). The *Δdum* mutant has a deletion of 175 bp within the second exon, and the resulting gene fusion is predicted, *in silico*, to encode a protein with the first 17 amino acids of tweedledum conserved before the breakpoint junction, after which a frameshift adds 7 missense amino acids followed by a stop codon (Supplementary Figure S3A–B). *Δdum* mutants engorged on blood meals to approximately double their body weight (Supplementary Figure S3C). They retained eggs and motile sperm in their visually healthy ovaries and spermathecae, respectively, for moderate (6 days post-blood-meal, Supplementary Figure S3D) or extended (12 days post-blood-meal, Supplementary Figure S3G) durations post-blood-meal. *Δdum* mutants laid a similar number of eggs compared to wild type after moderate retention (Supplementary Figure S3E-F), but significantly fewer eggs after extended retention (Supplementary Figure S3H-I). Of the eggs laid after extended retention, a starkly smaller proportion from *Δdum* mutants compared to wild type generated viable offspring (Supplementary Figure S3H,J). These results suggest that at minimum *tweedledum* is contributing to drought resilience, and future work will resolve if both *tweedledee* and *tweedledum* are required for this important phenomenon.

## DISCUSSION

### Flexibility enables a freshwater-centric lifestyle in variable environments

*Aedes aegypti* mosquitoes depend on freshwater availability for completing the aquatic larval and pupal stages of their life cycle (*20, 37, 39*). Adult *Aedes aegypti* females carrying mature eggs accordingly prefer to lay them at the edge of freshwater (*60*). Fluctuating climates with unpredictable and intense droughts likely impose selective pressures on this species, which has evolved multiple reproductive strategies that contribute to its resilience and invasive potential. Decoupling of mating from host-seeking and subsequent blood-feeding – such that either can take place first – and the appropriate coupling of both behaviors with egg-laying provides female *Aedes aegypti* mosquitoes with flexibility to maximize their reproductive output while still ensuring they have the required sperm and blood proteins for producing viable offspring. If faced with drought after being laid, embryos developmentally arrest in a state of diapause within the eggshell for several months as an added layer of protection, until freshwater that can support larval survival becomes available to stimulate hatching (*26*).

We suggest that the ability of adult *Aedes aegypti* females to retain mature eggs in their ovaries without complete loss of viability for flexible lengths of time while searching for an egg-laying site is of significant adaptive value. In this study, we demonstrated that *tweedledee* and *tweedledum*, a pair of linked, mosquito-specific genes, encode proteins that allow a female to retain her eggs for extended durations as needed without marked loss of viability, such as when access to freshwater is precluded due to drought. Females lacking both genes show a time-dependent phenotype. Their reproductive resilience dramatically worsens as length of egg retention increases from 6 to 12 days post-blood-meal, whereas wild-type females continue to maintain a remarkable degree of reproductive resilience regardless of the duration of egg retention (Figure 5J). Our findings highlight an example of plasticity in the innate reproductive behaviors of *Aedes aegypti* mosquitoes, which allows them to thrive in a remarkable range of ecosystems with distinct climates.

### Providing protection to eggs: abundance in the right place, at the right time

Oocytes are stored within the ovaries of females in species separated by millions of years of evolution. Increased oocyte storage time carries the risk of increased damage, which females have evolved diverse strategies to mitigate (*61*). Mammalian oocytes are maintained for decades and gradually released from their reserves. In humans, as a female and her oocyte reserve both age, oocytes become increasingly prone to meiotic segregation errors that result in higher rates of miscarriage and Down’s syndrome (*62*). Mammalian oocytes in reserve reside within primordial follicles where they are nurtured by maternally-derived nutrients that play a role in maintaining oocyte longevity (*62*). Bidirectional exchange between germline (oocyte) and somatic (follicle) cells is critical for germline maintenance and occurs through both gap junction-mediated transfer of small molecules, as well as via paracrine secretion of nourishing factors from follicles (*63*). In *Drosophila melanogaster* flies, oocytes are retained if access to protein or sperm is restricted, and extended oocyte storage results in lower capacity for embryonic development. In wild type fly oocytes, abundant expression of two heat shock protein chaperones, *Hsp26* and *Hsp27* contributes to maintenance of developmental capacity following extended oocyte retention (*61*). These examples highlight a strong precedence for the existence of protective or nurturing mechanisms in *Aedes aegypti* ovaries, which would enable the female to maintain viable eggs for extended durations post-maturation.

What is the mechanism by which a pair of abundantly expressed genes ensure long-term viability of eggs retained in the mosquito ovary? We have few clues to work with. The highly regulated spatiotemporal expression of these paired genes in the ovary suggests to us that they both contribute to egg viability during retention. After maturation, and in the hours post-egg-laying, expression of *tweedledum* is restricted within the ovaries to the follicular epithelial cells surrounding eggs. In contrast, *tweedledee* is basally expressed in all the reproductive states of the ovaries, although its upregulation by several orders of magnitude is concurrent with its expanded expression in the follicular epithelial cells surrounding mature eggs during egg retention, together with *tweedledum.* Since the follicular epithelial cells form a socket around mature eggs, the expression of both genes together in these cells may be most functionally relevant in protecting eggs during retention, such as by forming a secreted desiccation-resistant barrier or a coating required for long-term maintenance. On the way to being laid, mature eggs also interact with the calyx where *tweedledee* is additionally expressed. It is therefore possible that additional interaction between the eggs and *tweedledee* while eggs are in transit facilitates egg competence for subsequent sperm entry and fertilization after extended retention, thereby ensuring viability. Future biochemical studies may reveal mechanistic insight into whether the proteins interact with each other, or with other molecules that provide protection to the egg after maturation.

In this study, we generated both *Δdeedum* double mutants and a *Δdum* single mutant. Both strains have substantially similar phenotypes of reduced egg viability during extended retention. Our attempts to generate *Δdee* single mutants were unsuccessful. This leaves open the question of whether both *tweedledee* and *tweedledum* contribute to the *Δdeedum* phenotype or whether *tweedledum* is solely responsible.

### Signal, waste, or both?

What is the function of the circulating forms of these proteins? Our proteomic analysis supports the hypothesis that these proteins are secreted, as we detect tryptic peptides from both proteins that represent N-terminal signal peptide-cleaved forms, both in the circulating hemolymph, as well as in the ovary. It is unclear if the circulating forms of these proteins are waste products destined for destruction after production in the ovaries, or if they serve a signaling function. One possibility is that tweedledee and tweedledum sustain eggs in the ovary while also acting as a humoral signal to alert other organs – including the nervous system – of the female’s reproductive status. There is a precedent for secreted egg-related proteins serving multiple roles. Vitellogenins are best known as egg yolk proteins, but also function as a hormone secreted into hemolymph, where they have multiple effects on behavior and longevity in social insects (*64*). In *Aedes albopictus*, vitellogenins have recently been implicated in regulating host-seeking behavior in response to the status of nutritional reserves, in addition to their role as yolk protein precursors (*65*). Bifunctionality and coordinated action of molecular pairs are common phenomena in insect reproduction. The hormones, 20-hydroxyecdysone and juvenile hormone III, act in concert to control metamorphosis across insects (*66*), while additionally modulating vitellogenesis after a blood meal in mosquitoes (*44, 67, 68*) and defining caste-specific behavioral repertoires or traits in the ant, *Harpegnathos saltator* (*69*).

### Taxon-restricted genes underlie tailored adaptations in a diverse world

Adaptations, especially those relevant to reproduction or expansion into new ecological niches, have been shown in recent studies across diverse species to arise from taxon-restricted, rapidly evolving genes with tissue-restricted and/or sexually dimorphic expression (*70–72*). For example, a mouse *de novo* gene, *Gm13030*, with female-biased, oviduct-specific expression shows strong estrous cycle-dependent control (*73*). In homozygous *Gm13030* mutant females, three *Dcpp* genes known to promote embryo implantation are upregulated. Mutant females progress normally through their first estrous cycle but undergo premature implantation in their second estrous cycle, resulting in inappropriately early second litters and higher infanticide rates – both likely maladaptive phenotypes (*73*). In *Rhagovelia* water strider insects, a pair of taxon-restricted genes (*gsha* and *mogsha*) are required for the development of a midleg fan structure specific to this genus (*74*). The fan endows *Rhagovelia* with the biomechanical capabilities needed to perform a rowing behavior on the surface of rapidly moving streams where they are typically found (*74*). Other non-*Rhagovelia* species occupying the same streams rarely perform these rowing behaviors and instead occupy static surfaces on leaves, suggesting the fan is central to *Rhagovelia’s* ability to walk on water (*74*). *Hormaphis cornu* aphids secrete salivary gland-enriched BICYCLE proteins, which come from a large family of rapidly evolving secreted molecules. The aphids pierce their stylet into mesophyll cells of the witch hazel leaf, where they deposit BICYCLE proteins. This triggers the formation of galls on the witch hazel leaf, which provides the aphid with the shelter and nutrition required for subsequent development (*75*). *Aedes aegypti* and *Aedes albopictus* mosquitoes are both predicted to expand into new parts of the globe where they were previously absent, owing to their ecological flexibility (*76*). We propose that the taxon-restricted genes, *tweedledee* and *tweedledum*, are an integral part of the mechanisms allowing the ecological flexibility of this species without compromising on reproductive capacity.

### Evolutionary origins and catering to different natural histories

The evolutionary history of *tweedledee*, *tweedledum*, and the *conceptualogs* is intriguing. The lack of known domains poses a problem for understanding the function and evolutionary history of these proteins, and conventional homology searches fail to detect homologs that could provide clues. The two proteins have no homology to each other, to the conceptualogs, or to any other known protein. Structural homology-based approaches might be a path forward to identify genes with divergent sequence but conserved three-dimensional protein structure and function. However, current protein structure prediction programs perform poorly on small proteins, especially when phylogenetic homology cannot be brought to bear to inform the analysis (*77*).

Rapidly evolving genes typically show testis-biased expression across evolution (*72*), but in *Aedes aegypti*, ovary-specific genes evolve unusually fast with more frequently occurring signatures of positive selection as compared to genes with enriched expression in the testis (*78*). Our study reveals *tweedledee* and *tweedledum* as examples of such rapidly evolving, ovary-enriched genes present in mosquito genomes, but it remains an open question whether genes of this type exist outside of mosquitoes. Published transcriptomic data suggest the tissue-restricted and sexually dimorphic expression of these genes may be conserved across genera. In *Anopheles stephensi* (*79*), the *conceptualog* is upregulated in ovaries 24 hours post-blood-meal, and in *Anopheles arabiensis*, the *conceptualog* is upregulated in the reproductive tissues of females compared to males (*80*). At the genomic level, what allows the syntenic locus, characterized in several blood-feeding mosquitoes by the conserved *scratch* and *peritrophin-like* genes, to ‘trap’ one rapidly divergent gene in the case of *Culex* and *Anopheles* mosquitoes, or two rapidly divergent genes in the case of *Aedes* mosquitoes? Together, these observations of shared synteny and gene expression indicate that the *Aedes tweedledee/tweedledum*, and the *Culex* and *Anopheles conceptualogs* are likely to have evolved from a common ancestor, and that these genes may co-opt similar pathways across genera to function.

A look at the natural histories of different mosquito genera suggests that they each employ distinct life history strategies. These involve differences in adult female behavioral parameters: flexibility in choosing a blood meal host, circadian control of host-seeking, mating frequency, and egg-laying site selection; differences in the potential for diapause or resistance to desiccation in embryos; and tolerance of larvae for different aquatic environments (*24, 39*). The physiological and behavioral adaptations underpinning these strategies must co-evolve in each of the species, in turn determining the ecological niches that the species are able to exploit. Future comparative studies will resolve whether rapid divergence in the sequence of *tweedledee/tweedledum* and the *conceptualog* is accompanied by conservation, or by rapid divergence in their functions. This work thus highlights the importance of considering taxon-restricted genes as important points of study to understand the life-history strategies of a species, and to identify new inroads for breaking the cycle of mosquito-borne disease transmission.

## ACKNOWLEDGMENTS

We thank members of the Vosshall Lab for comments on the manuscript; Gloria Gordon and Libby Mejia for expert mosquito rearing; James Petrillo at The Rockefeller University Precision Instrumentation Technologies Resource Center for input on design, optimization, and fabrication of the metal blood puck; Vadim Sherman at the Rockefeller High-Energy Physics Machine Shop for fabrication of the metal blood puck; Rob A. Harrell II at the Insect Transgenesis Facility at the University of Maryland for CRISPR-Cas9 embryo injections; Benjamin J. Matthews for advice on the design of sgRNAs and egg-laying assays; Laura B. Duvall for advice on the design of host-seeking suppression behavioral assays; Katarzyna Cialowicz, Christina Pyrgaki, Carlos Rico, and Alison North at The Rockefeller University Bio-Imaging Resource Center for training and advice on confocal imaging (RRID:SCR_017791); Leah Houri-Zeevi for advice on RNA-seq experiment design, optimization, and data analysis; Thomas Carroll and Douglas Barrows for assistance with RNA-seq data analysis; Connie Zhao at The Rockefeller University Genomics Resource Center for assistance with RNA-seq library preparation and sequencing; Caroline Jiang for advice on statistical analysis of behavior data; Junhui Peng for discussion on protein structure predictions; Alexander Wild for mosquito photographs; Daniel Kronauer and Shai Shaham for general discussion and project guidance.

## FUNDING

Funding for this study was provided by the Boehringer Ingelheim Fonds Ph.D. fellowship (KV); a predoctoral fellowship from the Kavli Neural Systems Institute and NIH NIDCD grant F30DC017658 (MH); EMBO ALTF 286-2019 (NS); NIH MIRA R35GM133780, Robertson Foundation, Monique Weill-Caulier Career Scientist Award, Rita Allen Foundation Scholar Program, and a Vallee Scholar Program (VS-2020-35) (LZ); Fellowship of Tsinghua Xuetang Life Science Program (J. Zhao); NRSA Training Grant #GM066699 (LAN). The Proteomics Resource Center at The Rockefeller University acknowledges funding for mass spectrometer instrumentation from the Sohn Conferences Foundation and the Leona M. and Harry B. Helmsley Charitable Trust. LBV is an investigator of the Howard Hughes Medical Institute.

## AUTHOR CONTRIBUTIONS

KV carried out all experiments in this paper with the following exceptions: NS optimized and performed the ovary RNA-seq experiment for Figure 2 together with KV; PL conducted synteny, homology, whole-genome amino acid content, and dN/dS analysis for Figure 4 with guidance from LZ; SZ optimized protocols, with input from MH, and prepared all samples for RNA *in situ* hybridization of whole-mount ovaries for Figure 3; J. Zhao optimized protocols and together with KV conducted ovary proteomics for Figure 2 and provided practical assistance in generation of the *Δdeedum* strain for Figure 5. MH optimized protocols together with HM and performed hemolymph proteomics experiments with KV; J. Zeng designed and produced the blood puck assay; LAN provided practical assistance in conducting extended retention experiments to phenotype *Δdeedum* mutants in Figure 5. HM analyzed all proteomics data in Figure 2. LZ provided input on all evolutionary and genome architecture analyses in Figure 4. KV and LBV together conceived the study, designed the experiments and figures, discussed the results, and wrote the paper with input from all authors.

## DECLARATION OF INTERESTS

The authors declare no competing interests.

## MATERIALS AND METHODS

### Human and animal ethics statement

Behavioral experiments and blood-feeding methods using live hosts were approved and monitored by The Rockefeller University Institutional Review Board (IRB protocol LV-0652) and the Institutional Animal Care and Use Committee (IACUC protocol 17018), respectively. All human subjects gave their written informed consent to participate in this study.

### Mosquito rearing and maintenance

*Aedes aegypti* wild type (Liverpool) and CRISPR-Cas9 knockout strains were reared using standard insectary conditions in an environmental chamber maintained at 70-80% relative humidity and 25-28°C with a photoperiod of 14 hours light: 10 hours dark as previously described (*81*). Adults of all genotypes were provided *ad libitum* access to 10% sucrose and were housed in 30 cm^3^ BugDorm-1 Insect Rearing Cages (MegaView Science) unless otherwise specified. Newly generated mutant strains were blood-fed on human volunteers until they were established. For stock maintenance, females were blood-fed on live mice or on defibrinated sheep blood (Hemostat Laboratories, DSB100) using an artificial membrane feeder (the “blood puck”) described below. All animals used for behavior experiments, regardless of genotype, were blood-fed using the blood puck.

### Blood-feeding for behavior assays using the blood puck

For all behavior experiments requiring blood-fed mosquitoes, 5-16-day old females were fed sheep blood supplemented with 2 mM adenosine 5’-triphosphate (ATP) (Sigma Aldrich, A6419) in aqueous sodium bicarbonate buffer using a new artificial membrane feeder called the blood puck. Metal blood pucks were custom-made at The Rockefeller University Precision Instrumentation Technologies Resource Center and the Rockefeller High-Energy Physics Machine Shop. Three-dimensional designs for fabrication, and a bench manual for suggested use are provided (https://doi.org/10.5281/zenodo.5945525). The blood puck is a disc with one indented, rimmed face on which blood rests with Parafilm stretched over it. This allows the female mosquitoes to pierce the Parafilm membrane and feed on the blood beneath. The other face of the disc is fully flat and does not have Parafilm stretched across its surface.

Before assembling the blood puck, 8.1 mL of defibrinated sheep blood stored at 4°C was warmed to 42°C for 15-30 minutes in a water bath, and 1 mL aliquots of 20 mM ATP in 25 mM aqueous sodium bicarbonate stock stored at −20°C were slowly thawed on wet ice to room temperature. To assemble the feeding disc of the blood puck membrane-feeder, a 10×10 cm square of Parafilm M (Fisher Scientific, S37440) was first rubbed on both sides against a human skin surface free of cosmetics, such as the forearm or neck, then stretched evenly until translucent before setting aside. The blood puck disc was placed in a metal bead or water bath at 42°C for at least 10 minutes. It was then removed from the warming bath and thoroughly dried with a paper towel. Next, the Parafilm rubbed on human skin was stretched across the entire indented face of the disc with the utmost care taken to ensure there were no holes in the Parafilm on the feeding side of the disc. Additional strips of Parafilm were used to seal the edges of the disc, and the stretched Parafilm was checked to ensure that it was taut enough to be pierced by a female mosquito’s stylet. Working quickly to prevent heat dissipation from the pre-heated feeding disc and blood, 900 μL of ATP stock was added to the 8.1 mL of heated blood for a final concentration of 2 mM ATP, and vortexed thoroughly to mix. Care was taken to ensure the ATP was never heated and did not undergo multiple freeze-thaw cycles. The blood puck disc was held by its edges with the indented, rimmed side face-down and the flat side face-up. The blood + ATP mixture was pipetted through one of the two holes from the flat face. The disc was swirled laterally to evenly distribute the blood before gently placing the blood puck on top of a mesh face of the mosquito cage. In this configuration, the indented, rimmed side sat atop mesh of the cage with female mosquitoes beneath, while the flat side was face-up. Any excess blood dribbling out of the puck after placing on the mosquito cage was blotted with paper towels, and 1-2 additional metal discs (either additional blood pucks, or simple metal discs with both faces flat) pre-warmed to 42°C were placed on top of the feeding disc to maintain warmth. These discs were reheated and replaced as needed to maintain the feeding disc at an optimal temperature for mosquito blood-feeding. As needed, mosquitoes were activated by an experimenter exhaling their breath into the cage. Blood-feeding was conducted both at ambient room temperature conditions and in the environmental chamber with similarly high and reliable engorgement rates. Females were typically allowed to feed for 15 minutes, or until fully engorged, and a single blood puck could be used for 2-3 cages of ~400-450 females each with replacement of rewarmed flat discs between transfer of the apparatus between cages. After feeding to repletion, typically 15-30 minutes per cage, the discs were taken off the cage, the Parafilm discarded into biohazard waste, and the metal discs rinsed under hot water to thoroughly remove all traces of blood. The blood puck was dried with paper towels for subsequent use.

After feeding, animals were cold anesthetized in a 4°C cold room to separate and discard males, as well as non-blood-fed and partially engorged females. Fully engorged females were selected by eye and returned to their original rearing conditions in a fresh cage with continuous access to 10% sucrose.

### Preparation of mosquitoes for weighing

When blood meal size was measured by weighing (Figure 5C, Supplementary Figure S3C), non-blood-fed females of all genotypes were each split into two cages of 80-100 females and sugar-starved for 20-24 hours prior to delivering the blood meal. During the sugar-starvation period in experiments involving subsequent weighing, females were offered deionized water-soaked cotton balls to prevent dehydration. Non-blood-fed controls and experimental group females engorged on blood were both immediately cold anesthetized at 4°C after offering the blood meal and weighed in respective groups of 5 each, as previously described (*82*).

### Preparation of mosquitoes at different reproductive time-points

To prepare groups of mosquito females at different reproductive time-points, all groups were age-matched within each experiment to the extent possible and maintained in mixed-sex cages for at least 5-7 days post-eclosion to ensure that most females were mated. The only exception was with the “virgin” group (Figure 1B,F), for which females were separated at the pupal stage and maintained in single-sex cages. For all blood-feeding groups, females were provided sheep blood supplemented with 2 mM ATP, and only fully engorged females were selected by eye for subsequent experimental use. For egg retention groups, cages were carefully checked for any prematurely dumped eggs prior to collection of females for dissections. For all experiments in which females were required as soon after egg-laying as possible, i.e., groups where eggs were laid <5 hours prior (Supplementary Figure S1C–D, Figure 2, and Figure 3), we standardized 3 hours as the allotted time for individual females after they were aspirated into egg-laying vials at room temperature. The allotted time of 3 hours was determined based on our finding that ~80% of females complete egg-laying within 3 hours of transfer to egg-laying vials (Supplementary Figure S1A–B). Eggs laid by *Aedes aegypti* females are initially white, and melanize within the first 1-2 hours of egg-laying (*52*). Based on this, we postulated that any egg-laying vials with at least 10 melanized eggs are likely to have come from females that had completed laying their full clutch of ~100 eggs.

Females of the <5 hours post-egg-laying group used for ovary RNA-sequencing, ovary proteomics, and hemolymph proteomics experiments in Figure 2 were tested behaviorally to verify that they had restored attraction to humans using a long-range, live human stimulus olfactometer as described (*83*). Briefly, females that had laid ≥10 melanized eggs were pooled into groups of 20, gently aspirated at room temperature into “start” canisters of the olfactometer, acclimated for 10 minutes, and the trial run for 5 minutes and 30 seconds as per standard assay protocol. Attracted females were defined as those that entered the trap proximal to the human arm. These attracted females were collected directly from the trap, and only these attracted females were dissected for ovary or hemolymph sample collection. Females that did not enter the attraction trap were discarded and not used for ovary or hemolymph sample collection.

For whole-mount ovary fluorescence RNA *in situ* hybridization experiments in Figure 3B–E, females were collected and used immediately from egg-laying vials that contained at least 10 melanized eggs, without further assessment of their attraction to humans. For groups that had laid eggs greater than 1 week prior to sample collection (Figure 3F), plastic cups (VWR HDPE Multipurpose Containers, H9009-664) half-filled with deionized water and lined with filter paper (GE Healthcare, WHA1001055) were introduced to the cage as continuously available egg-laying substrates between 3 and 6 days after blood-feeding. When dissected, 13 days had elapsed since the last blood-meal of this group. When groups in their second reproductive cycle were collected, either for behavior (Figure 1E) or for hemolymph proteomics (Figure 2L–N), they were treated equivalently to the corresponding first reproductive cycle groups.

### Live human olfactometer assay

Live human olfactometer assays for testing female mosquito attraction to a human forearm were performed as previously described (*83*). The same subject was used as a stimulus in all experiments. Fabrication and assembly details, as well as a user guide are available at https://github.com/VosshallLab/Basrur_Vosshall2020.

Experiments were conducted at 70-90% humidity and 25-28°C. Each trial consisted of approximately 20 female mosquitoes, grouped by reproductive condition. The groups of 20 were aspirated into “start” canisters at least 30 minutes before their trial. Females were given continuous access to 10% sucrose prior to sorting into canisters but were not provided any sucrose or water after being re-housed in the canister. Trials with non-blood-fed females were treated as positive controls and were interspersed throughout each experimental day. Experimental days were counted for final analysis only after ensuring average attraction to the live human arm stimulus of the non-blood-fed group was ≥50% across trials. All groups were run on each experimental day. Two trials were run simultaneously, one using each arm of the live experimenter as the stimulus. Groups were shuffled between ports to minimize bias.

### Egg retention, laying, and hatching assay

For all egg retention experiments, to prevent accumulated condensation that could trigger premature egg ‘dumping’, special care was taken to ensure that all cages and sucrose-soaked cotton wicks for blood-fed female mosquitoes were not subjected to frequent fluctuating temperature and humidity, and closely monitored to remove any accumulated droplets of water. Following the duration of egg retention, cages were thoroughly checked for any dumped eggs. If a small proportion of eggs was found prematurely dumped on the sugar-soaked cotton wicks, this was noted prior to set up of egg-laying. Although observed extremely rarely, if the cage floor was found to be covered with a large proportion of prematurely dumped eggs, suggesting that little to no egg retention was achieved, all females in such a cage were discarded and not used for further experimentation.

Only for the data shown in Supplementary Figure S2D, a single-female modular egg-laying assay setup was used as described (*21*). The assay setup was modified to accommodate 28 females instead of 14, and each female was provided access to a single egg-laying substrate of deionized water instead of two substrate choices. Details of design and fabrication are available at https://github.com/VosshallLab/MatthewsYoungerVosshall2018.

For all other egg-laying behavior experiments, at the time of egg-laying, females with retained eggs were 14-21 days old, and aspirated out of their cages at room temperature into individual egg-laying vials. Egg-laying vials were made using plastic *Drosophila* vials (VWR, 25 mm diameter, 95 mm length, 75813-164) with 2-3 mL of deionized water, and a 55 mm diameter Whatman filter paper (GE Healthcare, WHA1001055) folded into a cone at the bottom of the vial to serve as a moist egg-laying substrate as previously described (*21, 35*). Vials were kept plugged (Genesee Scientific, Flugs® Narrow Plastic Vials, 49-102) under standard insectary conditions following transfer of females ready for egg-laying. All females were removed 20-24 hours after transfer under brief cold anesthesia and either discarded or stored at –20°C if required for further genotyping. Filter paper lined with laid eggs, and any eggs remaining on the sides of the vial or in the water, were removed from the vial at room temperature and placed briefly on a paper towel to remove excess moisture. The eggs were then manually counted by eye or under a dissection scope as needed and the number of eggs laid per female recorded. If most eggs from a female were unmelanized or submerged in the pool of water instead of lined on the filter paper, the sample was excluded. The egg-lined filter paper was returned to the emptied and dried vial within 24 hours of removing the female and terminating the egg-laying assay. All vials were kept under standard insectary conditions for 6-14 days prior to hatching.

Egg hatching was staggered to ensure that all egg papers were dried for the same length of time prior to hatching (6-14 days), and egg papers from distinct individuals were hatched and maintained separately. Eggs were hatched either by transferring egg papers into a small plastic cup (VWR, HDPE Multipurpose Containers, H9009-662) with 50-60 mL ‘hatch broth’ comprised of deoxygenated water with finely ground fish food (Pet Mountain, Tetramin Tropical Tablets Fish Food for Bottom Feeders, YT16110M), or by adding 20 mL of hatch broth directly to the egg-laying vial with the dry egg paper (Figure 1, and Figure 5 or Supplementary Figure S3, respectively). Larvae hatched were provided with a fresh pinch of fish food as needed.

At least 5 days after hatching, the egg viability experiment was terminated. Egg papers were removed and larvae, sometimes mixed with pupae or eclosed adults, were either cold anesthetized at 4°C or killed by freezing at – 20°C overnight before thawing and counting. Offspring from each individual female were separately poured onto Petri plates (Fisher Scientific, S33580A) or small plastic cups with fresh deionized water and photographed on a light board using a webcam (Logitech, C922x Pro Stream Webcam) mounted from above, ensuring that all offspring were captured in the field-of-view. Captured images were imported into FIJI/ImageJ (NIH), and the number of offspring from each individual female was counted using the Cell Counter plugin.

### Bulk RNA-sequencing of mosquito ovaries

For ovary bulk RNA-sequencing (RNA-seq), 3 pairs of ovaries were used for each replicate, and 4 replicates were prepared per experimental group from 19-to-20-day old females. Mosquitoes were cold-anesthetized and kept on ice for up to 1 hour, or until dissections were complete. Ovaries were dissected on ice, in ice-cold RNase-free 1X phosphate-buffered saline (PBS) (Invitrogen, AM9625). They were moved using forceps into 0.5 mL Eppendorf LoBind microcentrifuge tubes (Sigma Aldrich, Z666521), and immediately snap-frozen on a cold block (Simport, S700-14) pre-chilled to −78°C on dry ice. Extreme caution was taken during the tissue dissection to ensure that there was no contamination from other mosquito tissues. Each dish and forcep was carefully cleaned with 70% ethanol and RNase-away (Thermo Fisher, 7003) after every dissection. All replicates for each experimental group were dissected in parallel to avoid artifacts and batch effects. Dissected tissue was stored at −80°C until RNA extraction.

RNA extraction was performed using the PicoPure Kit (Thermo Fisher, KIT0204) with the following modification for homogenizing tissue: instead of lysis buffer, 100 μL of TRIzol (Thermo Fisher, 15596026) was added to the collection tube on ice. Tissues were homogenized manually using a Pellet Pestle Motor (Kimble, 749540) and an RNase-Free pellet pestle (VWR, KT749510-0590) for 30 seconds following the addition of 140 μL of TRIzol to a total of 240 μL. Tubes stood at room temperature for 5 minutes before 48 μL of chloroform:isoamyl alcohol 24:1 was added (Sigma, C0549). Tubes were hand-shaken for 30 seconds and left to stand for 2 minutes before centrifuging at 12,000 RPM for 15 minutes at 4°C. The aqueous TRIzol layer was then removed and added into the PicoPure column, up to 130 μL at one time. Subsequent steps were performed according to PicoPure manufacturer’s instructions, including DNase treatment.

Samples were run on a Bioanalyzer RNA Pico Chip (Agilent, 5067-1513) to determine RNA quantity and quality. RNA quantity was re-verified with a Qubit 2.0 Fluorometer using the RNA HS Assay Kit (Invitrogen, Q32855). The three biological replicates with the most consistent RNA yield across conditions were then used for library preparation and sequencing.

100 ng of total RNA was used to generate RNA-seq libraries using Illumina TruSeq stranded mRNA LT kit (Illumina, 20020594), following the manufacturer’s protocol. Libraries prepared with unique dual indexes were pooled at equal molar ratios. Sequencing was performed at The Rockefeller University Genomics Resource Center on the Illumina NovaSeq 6000 sequencer using V1.5 reagents, the SP flow cell, and NovaSeq Control Software V1.7.0 to generate 150 bp paired end reads, following manufacturer protocol. Data were demultiplexed and delivered as fastq files for each library. Sequencing reads have been deposited at the National Center for Biotechnology Information (NCBI) Sequence Read Archive (SRA) under BioProject PRJNA796320. The data discussed in this publication have been deposited in NCBI’s Gene Expression Omnibus (*84*) and are accessible through GEO series accession number GSE193470 (https://www.ncbi.nlm.nih.gov/geo/query/acc.cgi?acc=GSE193470).

### Alignment and quantification of RNA-seq data

Sequence and transcript coordinates for the *Aedes aegypti* mosquito genome and gene models were obtained by merging the Aaeg_L5 RefSeq annotation from NCBI with a manual chemoreceptor annotation. Information related to generating this annotation is available at https://github.com/VosshallLab/Jove_Vosshall_2020/tree/master/RNAseq_merged_annotation (*85*). Transcript expression was calculated using the Salmon quantification software (version 0.8.2) (*86*), and gene expression levels as transcripts per million (TPMs) and counts were retrieved using Tximport (version 1.8.0) (*87, 88*). A table of TPM counts for all reproductive conditions and replicates can be found on Zenodo (https://doi.org/10.5281/zenodo.5945525). Normalization and rlog transformation of raw read counts in genes were performed using DESeq2 (version 1.20.0) (*89*). The normalized and transformed counts were used to perform principal component analysis (PCA) using DESeq2, and to assess between-sample variability with hierarchical clustering and with calculation of sample distance correlations (*89*).

### Ovary collection and sample preparation for proteomics

To extract whole proteins from ovaries, 8 pairs of ovaries were used for each replicate, and 4 replicates were prepared per experimental group. Ovaries were dissected in a droplet of 1X PBS, as needed, and boiled for 5 minutes at 100°C in 150 μL MilliQ water. Samples were centrifuged at 12,000 RPM for 30 seconds. The water fraction was then decanted into a separate tube and set aside. Extraction solution (150 μL of 0.25% acetic acid) was added to the precipitate, and the tissue was homogenized with a 5 mm tungsten carbide bead in a bead mill homogenizer (Qiagen, Tissue Lyser II) at 30 Hz for 1.5 minutes. The water and acid fractions were centrifuged separately at 4°C, 8000 RPM for 30 minutes. The two supernatants were then combined and spun to dryness in an Eppendorf Speedvac at 60°C for 1-1.5 hours. The mass spectrometry proteomics data have been deposited to the ProteomeXchange Consortium via the PRIDE (*90*) partner repository with the dataset identifier PXD030925. Ovary sample raw files begin with the code “MS205850LUM”.

### Hemolymph collection and sample preparation for proteomics

To collect hemolymph, 5 females were used per replicate. Cold anesthetized females were kept on ice and decapitated using 2.5-mm cutting edge Vannas spring scissors (Fine Science Tools, 15000-08) under a dissection microscope at 10X. Cold 30 μL1X PBS with 0.05% Tween (PBS-T) was pipetted as a bubble onto a 35 mm Petri plate (Falcon, 351008) on ice. The decapitated thorax was positioned close to the droplet without touching, and the thorax was gently squeezed using blunt forceps to release hemolymph from the decapitation site into the droplet of PBS-T. This was repeated such that each droplet of PBS-T consisted of pooled hemolymph from a total of 5 females for a single replicate. The PBS-T with hemolymph was pipetted into a 1.5 mL Eppendorf Protein LoBind tube (Thermo Fisher), and the Petri dish was washed with 10 μL PBS-T, and the 10 μL wash was combined with the ~30 μL from the initial extract. The samples were then heat-inactivated at 90°C for 10 minutes, snap-frozen on dry ice and stored in Eppendorf Protein LoBind tubes at −80°C until the subsequent steps could be carried out. For acetone precipitation of the extracted proteins, we ensured all samples, reagents, tubes, and tube racks were maintained at −20°C. Hemolymph samples from −80°C were quickly removed onto racks cooled to −20°C after which 6 volumes of acetone (~210 μL) cooled to −20°C were added. The sample tubes were vortexed for a few seconds until the frozen hemolymph samples fragmented and mixed well with the acetone. The tubes were then incubated upright at −20°C overnight. Following incubation, samples were spun down at 13,000 x g at 4°C for 10 minutes in a tabletop microcentrifuge. Most of the supernatant was removed with a pipette and discarded, leaving the protein pellet wet before storing at −80°C until subsequent steps. The mass spectrometry proteomics data have been deposited to the ProteomeXchange Consortium via the PRIDE (*90*) partner repository with the dataset identifier PXD030925. Hemolymph sample raw files begin with the code “MS195106LUM”.

### Liquid chromatography-mass spectrometry (LC-MS)

Dry protein pellets of both ovary and hemolymph samples were dissolved and reduced in 8 M urea (Fisher Scientific, 45000234)/70 mM ammonium bicarbonate (Fisher Scientific, 501656826)/20 mM dithiothreitol (Sigma Aldrich, 233153), followed by alkylation in the dark with 50 mM iodoacetamide (Sigma Aldrich, I1149). Samples were then diluted 2-fold and digested overnight with endoproteinase LysC (Fujifilm Wako Chemicals, WAKA Lysyl Endopeptidase, 129-02541). Samples were additionally diluted 2-fold and digested with trypsin (Promega, Sequencing Grade Modified Trypsin, Lyophil, PRV5111) for 6 hours. Digestions were halted by acidification and peptides were solid phase-extracted prior to analysis by LC-MS/MS. Peptide samples were analyzed by nano-flow LC-MS/MS (EasyLC 1200) coupled to a Fusion Lumos (Thermo Fisher) operated in High/High Data Dependent Acquisition (DDA) mode using Lock mass m/z 445.12003. Peptides were separated by reversed phase chromatography using 12 cm/75 μm, 3 μm C 18 beads (Nikkyo Technologies, NTCC-360/75-3-123°Column) with buffer A: 0.1% formic acid (Fisher Scientific, A11750), and buffer B: 80% acetonitrile (Fisher Scientific, A955) in 0.1% formic acid. For the hemolymph samples, a gradient from 2% buffer B/98% buffer A to 35% buffer B/65% buffer A in 70 minutes was used. For the ovary samples, a gradient from 2% buffer B/98% buffer A to 38% buffer B/62% buffer A in 90 minutes was used.

Data were queried against ‘GCF_002204515.2_AaegL5.0_protein.fasta’ database using MaxQuant software with the Andromeda search engine v.1.6. 6.0 (*91*). Oxidation of methionine and N-terminal protein acetylation were allowed as variables, and cysteine carbamidomethylation was defined as a fixed modification. Mass tolerance was set at 4.5 parts per million (ppm) for precursor ions and 20 ppm for fragment ions. Two missed cleavages were allowed for specific tryptic database searches. The ‘match between runs’ setting was enabled. False discovery rate (FDR) for proteins was set at 1% combined with a peptide FDR of 2%. Intensity based absolute quantitation (iBAQ) (*92*) values were used as a proxy for protein abundances. Data were processed using Perseus <u>v.1.6.10.50 (*93*)</u>. Reverse database hits and contaminating proteins were removed, and it was required that a protein was to be measured (using iBAQ) in at least 3 of 4 replicates for least one of the experimental groups. Each log^2^-transformed iBAQ signal was normalized by subtracting the respective sample’s median iBAQ signal. Missing values were assumed ‘Missing Not At Random’ (MNAR) (*94*) and a random distribution of signals with a width of 0.3 and a downshift of 1.8 were used to impute missing values. The sample sets were assessed for quality and correlation using scatter plots and PCA. Tables of iBAQ values and other analyzed metrics are available on Zenodo (https://doi.org/10.5281/zenodo.5945525) for all reproductive conditions and replicates for both ovary and hemolymph proteomics datasets.

### Whole-mount ovary fluorescence RNA *in situ* hybridization

The previously described hybridization chain reaction (HCR) technique (*95, 96*) was modified to detect RNA in whole-mount ovaries. All reagents, including custom probes, amplifiers, Probe Hybridization Buffer, Amplification Buffer, and Probe Wash Buffer, were purchased from Molecular Instruments. Adult female mosquitoes were dissected ~20 days post-eclosion. They were grouped by reproductive condition, cold anesthetized at 4°C, and maintained on ice for 30 minutes while ovaries were dissected from each female in 0.1X PBS. Dissected ovaries were incubated in a solution of 4% paraformaldehyde, 1X PBS, and 0.03% Triton X-100, and rotated overnight at 4°C. Ovaries were then washed 4 times in 1X PBS containing 0.1% Tween-20 (0.1% PBS-T) for 10 minutes each. Subsequently, ovary samples were dehydrated on ice using a series of graded methanol/0.1% PBS-T washes for 10 minutes, as follows: 25% methanol in 0.1% PBS-T, 50% methanol in 0.1% PBS-T, 75% methanol in 0.1% PBS-T, and two washes in 100% methanol. Ovaries remained in 100% methanol at −20°C overnight. To rehydrate the ovaries, samples were washed for 10 minutes each on wet ice with a series of graded methanol/0.1% PBS-T solutions, as follows: 75% methanol in 0.1% PBS-T, 50% methanol in 0.1% PBS-T, 25% methanol in 0.1% PBS-T, and two washes of 0.1% PBS-T. Ovary tissue was then digested in 20 μg/mL Proteinase-K (Thermo Fisher, AM2548) with 0.1% PBS-T for 30 minutes at room temperature and subsequently washed twice in 0.1% PBS-T at room temperature for 10 minutes each. Tissues were then fixed in 4% paraformaldehyde in 0.1% PBS-T for 20 minutes at room temperature and washed 3 times in 0.1% PBS-T for 15 minutes each at room temperature.

Ovaries were incubated in Probe Hybridization Buffer for 5 minutes at room temperature, and subsequently in a 37°C hybridization oven for 30 minutes in pre-warmed Probe Hybridization Buffer. A solution of pre-warmed Probe Hybridization Buffer and probe sets, each at 8 μmol, was mixed, and used to incubate samples at 37°C in a hybridization oven for three nights. Ovaries were next washed 5 times for 10 minutes each in a 37°C hybridization oven using Probe Wash Buffer pre-warmed to 37°C. The samples were then washed twice with 5X saline-sodium citrate (SSC) buffer (Invitrogen, 15557044) containing 0.1% Tween-20 solution for 5 minutes each at room temperature. To pre-amplify, ovaries were incubated in room temperature Amplification Buffer for 10 minutes. 24 μmol hairpins were prepared by heating 8 μL of 3 μM stock of H1 and H2 hairpins, separately, each at 95°C for 90 seconds on an Eppendorf Mastercycler. The hairpins were cooled to room temperature for 30 minutes in the dark, as hairpins are photosensitive and subject to photobleaching. Hairpins were then added to 100 μL of Amplification Buffer in which ovaries were incubated on a rotator at room temperature in the dark overnight. Ovaries were next incubated in the dark in a solution of 1:1000 DAPI in 5X SSC with 0.1% Tween-20 at room temperature for one hour. Ovaries were finally washed four times for 10 minutes each in 5X SSC with 0.1% Tween-20 and mounted in SlowFade Diamond (Thermo Fisher, S36972) onto glass slides with confocal microscopy-compatible coverslips.

### Ovary confocal imaging

Ovary images were acquired using an Inverted LSM 880 Airyscan NLO laser scanning confocal and multiphoton microscope (Zeiss). Either a 10x/0.45 NA objective, or an immersion-corrected 25x/0.8 NA or 63x/1.4 NA objective was used at a resolution of 1024 x 1024 pixels. If tiling was used, images were stitched with 10% or 12% overlap. Laser power, gain, and other parameters were individually optimized to acquire highest quality images for ovary samples acquired from non-blood-fed and post-egg-laying animals. Confocal images were viewed and processed using FIJI/ImageJ, and single slices were selected as representative images.

### Identification or orthologs and conceptualogs

Orthologs for *tweedledee* and *tweedledum* in *Aedes albopictus* were identified using orthology relationships in VectorBase. We noted that the current release of the *Aedes albopictus* genome assembly (GCF_006496715.1 as of December 2021) contained multiples copies of the locus with *tweedledee*, *tweedledum*, *scratch* and *peritrophin-like*. With the currently available data, we were unable to ascertain whether the multiple copies of the locus reflect true duplication events, or incompletely collapsed haplotypes. We arbitrarily chose one locus for subsequent analyses with *Aedes albopictus* genes, but repeating analyses with genes in a second locus yielded no significant differences. *Conceptualogs* in *Culex quinquefasciatus* and all specified *Anopheles* species were found by searching syntenic genomic regions between annotated orthologs of *peritrophin-like* and *scratch* on VectorBase. *Anopheles* genome annotations were used for further analyses if *peritrophin-like, scratch* exon 1 and the *conceptualogs* were all unambiguously annotated, and if all 3 genes were found on the same contig. *Aedes* and *Culex* protein sequences were obtained from NCBI. *Anopheles* protein sequences were also downloaded from NCBI if a RefSeq annotation was available, but if unavailable as with *Anopheles quadriannulatus* and *Anopheles culicifacies*, protein sequences were taken from VectorBase. Multiple sequence alignments and protein sequence identity matrices were generated using MUSCLE (*97*). The annotation of *scratch* was fragmented, with both exons annotated as separate genes in *Aedes aegypti*, *Anopheles arabiensis*, *Anopheles culicifacies* and *Anopheles quadriannulatus*, likely due to the presence of a large >50 kb intron. To minimize the ambiguity of the *scratch* protein sequences in these species, we generated the multiple sequence alignment between only the first exon of *scratch* in each species. Gene accession numbers are available at https://doi.org/10.5281/zenodo.5945525.

### Guanine+cytosine (GC) content analysis

GC content for all protein-coding genes from *Aedes aegypti* (AaegL5) was retrieved from Ensembl Metazoa BioMart (version 0.7) (*98*) using VectorBase as the gene source. The search was then limited to genes with a predicted cleavage site (SignalP 4.1) to filter for protein-coding genes with a predicted signal peptide.

### Amino acid content analysis

All protein sequences with a predicted signal peptide encoded by the *Aedes aegypti* (AaegL5) genome were retrieved from Ensembl Metazoa BioMart (version 0.7) (*98*). The signal peptide predicted was cleaved for each protein using SignalP 4.1 (*99*), and the percent of each amino acid was then calculated for the cleaved sequence. Mean percent residue was calculated for 3,040 *Aedes aegypti* proteins (minimum protein length = 60 amino acids) with predicted signal peptides.

### dN/dS ratio

We aligned coding sequences of 8,030 protein-coding *Aedes aegypti* genes to unique orthologs in *Aedes albopictus*, and coding sequences of 9958 protein-coding *Anopheles gambiae* genes to unique orthologs in *Anopheles stephensi*, as annotated in Ensembl Metazoa, via PRANK (*100*) using the codon option. dN/dS values per gene were calculated with KaKs_calculator (*101*) using the YN model (*102*). Sliding window values of dN/dS for *Aedes tweedledee* and *tweedledum* were calculated using a custom script for KaKs_calculator available at https://github.com/LiZhaoLab/Kaks_Calculator.

### Generation of *Δdeedum* double mutants

The *Δdeedum* double mutant was generated using CRISPR-Cas9 (*103*). Wild type embryos of the *Aedes aegypti* Liverpool strain were injected at the Insect Transformation Facility at the University of Maryland Institute for Bioscience and Biotechnology Research with a gene-targeting mixture composed of 300 ng/μL Cas9 protein with NLS (PNA Bio, CP01-200) and 4 sgRNAs, each 40 ng/μL. Two of the sgRNAs targeted exon 2 of *tweedledee* and the other two targeted exon 2 of *tweedledum* (see https://doi.org/10.5281/zenodo.5945525). Coordinates on *Aedes aegypti* chromosome 2 for *tweedledee*: 113,795,266 - 113,794,685 and *tweedledum:* 113,807,172 - 113,806,119, as annotated in the AaegL5 genome (*40*). As described (*103*), DNA templates were generated for each sgRNA by annealing oligonucleotides using the NEBNext Ultra II Q5 master mix (NEB, M0544L). The HiScribe Quick T7 kit (NEB, E2050S) was then used for in vitro transcription, per manufacturer’s instructions, with an overnight incubation of 17 hours at 37°C. Prior to mixing with Cas9 protein, sgRNAs were purified using SPRI beads (Beckman-Coulter, Ampure RNAclean, A63987) with elution in Ultrapure DNase/RNase-free distilled water (Invitrogen, 10977-015). The mutant allele was identified using polymerase chain reaction (PCR) and confirmed to be a double mutant, *Δdeedum*, in which both *tweedledee* and *tweedledum* were disrupted. The strain was backcrossed to wild type Liverpool animals for a minimum of four generations before inbreeding to homozygose. The homozygous *Δdeedum* strain was successfully established and behaviorally phenotyped. To verify wild type, +/*Δdeedum*, and *Δdeedum*/*Δdeedum* animals, 3 independent PCRs were run on each of the DNA template genotypes as described in Supplementary Figure S2A. The corresponding genotyping primers are listed at https://doi.org/10.5281/zenodo.5945525.

### Generation of *Δdum* single mutant and attempted generation of *Δdee* single mutant

*Δdum*, a *tweedledum* single mutant that was wild type at the *tweedledee* locus was recovered using the same mutagenesis procedure as the *Δdeedum* double mutant described above. The same genotyping strategy as that used for the *Δdeedum* double mutant was used with the *Δdum* single mutant to confirm that only *tweedledum* was mutated. With a distinct cocktail of sgRNAs, an allele with a deletion in *tweedledee* (*Δdee*) that spared *tweedledum* was also isolated but attempts to homozygose and establish a *Δdee* mutant have been unsuccessful to date.

### Photographs of ovaries and spermathecae

Mosquitoes were cold-anesthetized and kept on ice for up to 1 hour, or until dissections were complete. Ovaries and spermathecae were dissected on ice in 1X PBS. Ovaries were photographed using an AxioCam ERc 5s camera (Zeiss) attached to a stereo microscope (Zeiss, SteREO Discovery KMAT). Spermathecae were photographed using an iPhone X through the iDu Optics® LabCam® adapter attached to the eyepiece of a wide-field compound microscope (Swift, SW350B).

### Statistical analysis

R (version 4.1.1) and GraphPad Prism 9 software were used for data visualization and statistical analysis except when specified otherwise in the sections above or in the figure legends.

## DATA AND RESOURCE AVAILABILITY

All raw data reported here, along with a TPM count table from ovary RNA-seq, two protein abundance (iBAQ) tables from hemolymph and ovary proteomics respectively, and instructions for fabricating and using the blood puck are available on Zenodo (https://doi.org/10.5281/zenodo.5945525).

---

**Supplementary Figure S1.**
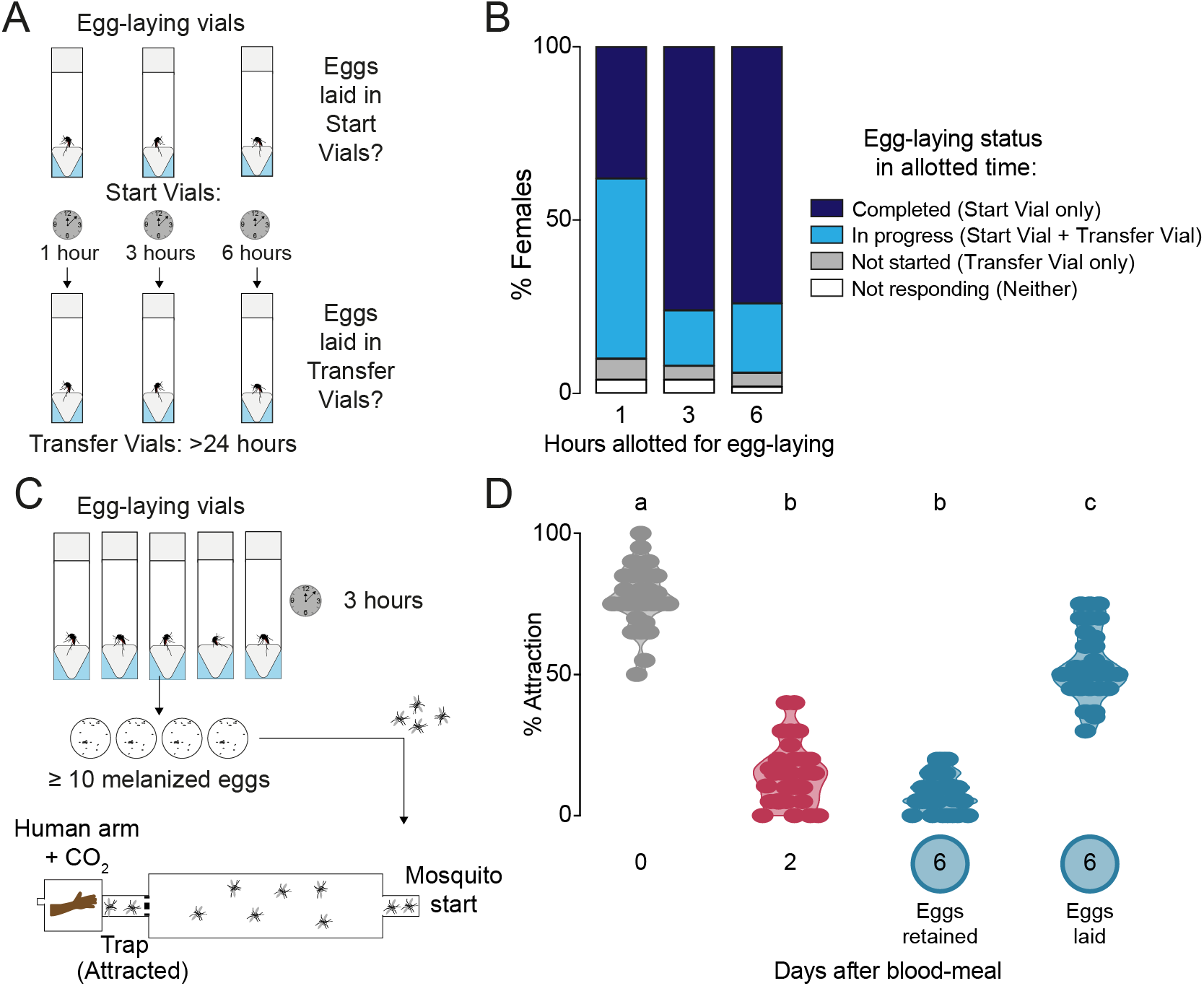
– related to Figures 1–3. Timing of egg-laying and return to human host seeking. **(A)** Assay schematic to determine the length of time taken for individual wild-type females to complete egg laying. **(B)** Temporal dynamics of egg-laying (n=50 females/time-point) according to criteria indicated in legend at right. **(C)** Cartoon of independent assay to verify and collect females that have restored attraction to humans as soon after egg-laying as possible for ovary RNA-sequencing, ovary proteomics, and hemolymph proteomics (Figure 2). Females were given 3 hours for egg-laying based on results in (B) and only females that had laid at least 10 melanized eggs – those presumed to have completed egg-laying – were chosen for subsequent testing of attraction to humans within an additional 2 hours using a long-range, live human stimulus olfactometer (*83*). **(D)** Attraction of wild type females to a human forearm at indicated reproductive state. Violin plot with median and 1st/3rd quartiles and showing all data points. Each point represents a single trial with a group of ~20 female mosquitoes with n=30-31 replicates/group. Data labeled with different letters are significantly different (one-way ANOVA, Tukey’s multiple comparisons test, p<0.0001).

**Supplementary Figure S2.**
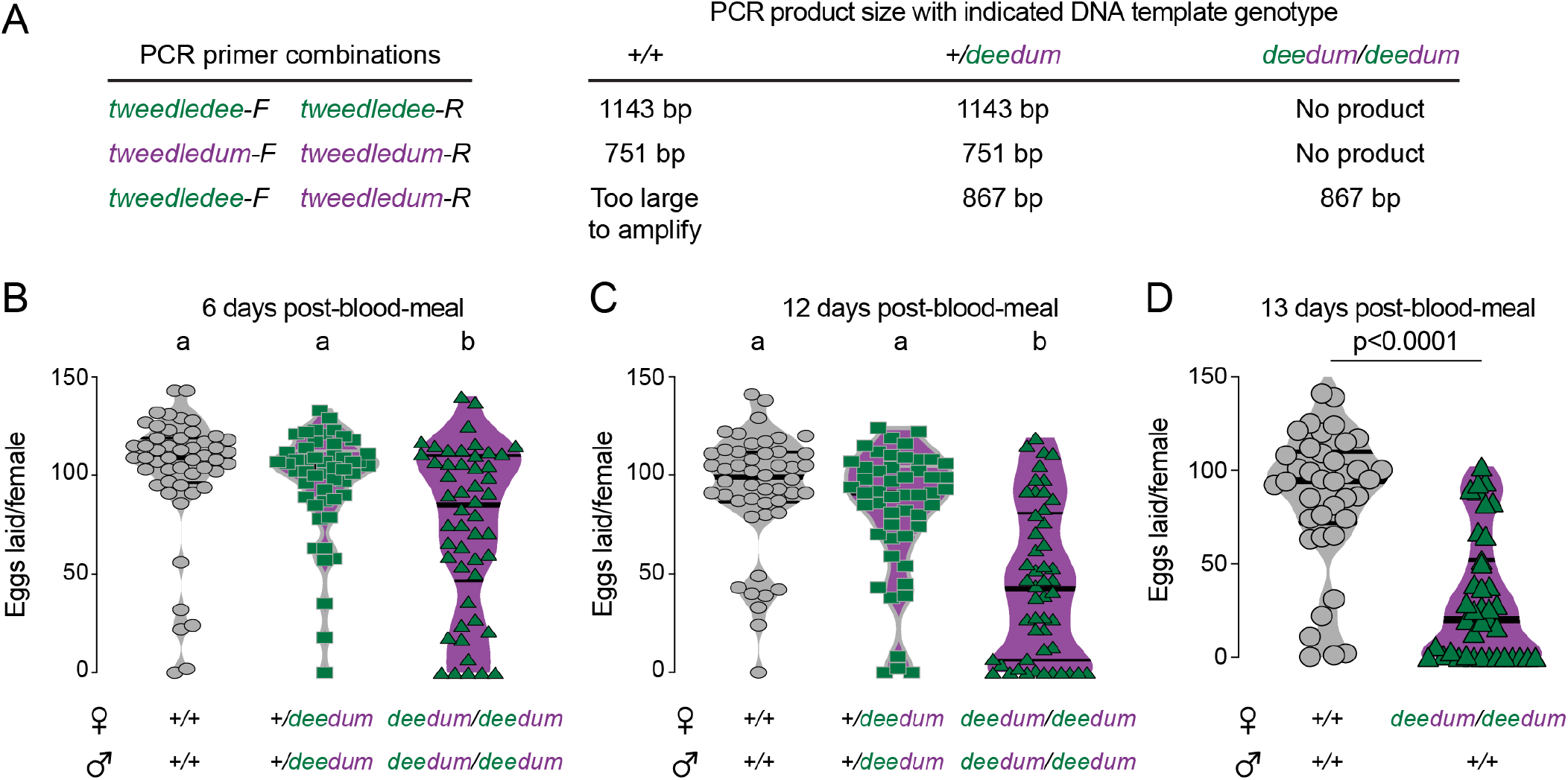
– related to Figure 5. *Δdeedum* double mutant genotyping and additional egg-laying data. (**A**) Genotyping strategy for *Δdeedum* double mutants. Primer sequences are at https://doi.org/10.5281/zenodo.5945525. (**B-D**) Eggs laid by females of the indicated genotype 6 (B), 12 (C), or 13 (D) days post-blood-meal. The genotype of the males to which they were mated is indicated below the female genotypes. Data are plotted as violin plots with median and 1st/3rd quartiles and showing all data points. Each point represents the eggs laid by a single female (B: n=50/genotype; C: n=48-50/genotype; D: n=38-39/genotype). In B,C, data labeled with different letters are significantly different (Kruskal-Wallis, Dunn’s multiple comparisons test, p<0.05). In D significance established by Mann-Whitney test, p<0.0001).

**Supplementary Figure S3.**
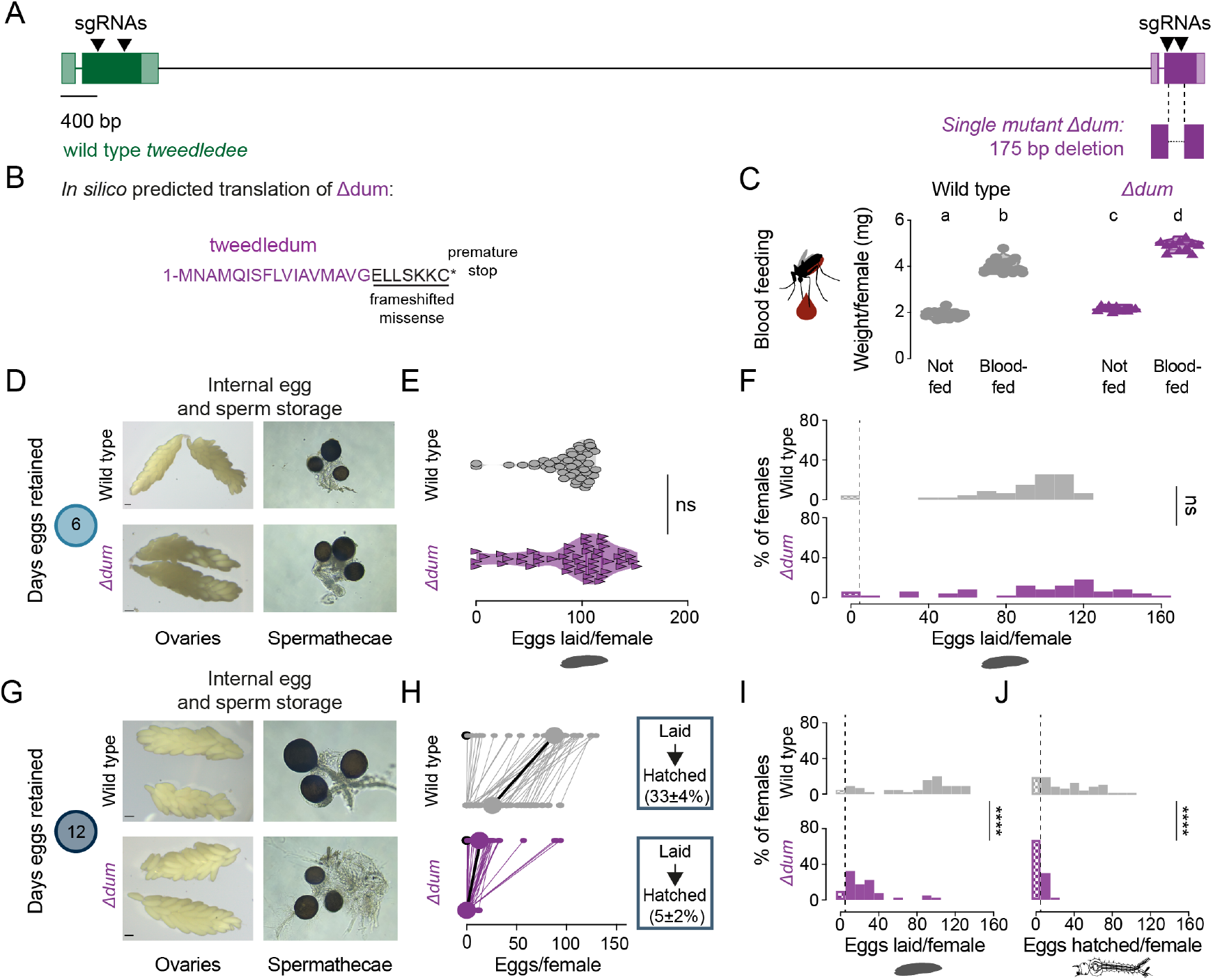
*tweedledum* mutants show defects in reproductive resilience during drought. **(A)** Schematic of *Δdum* mutant that deletes *tweedledum*, but not *tweedledee*. **(B)** *In silico* predicted translation of Δdum. **(C)** Averaged weights of 5 females of the indicated genotype not fed or blood fed, n=11-16 groups of 5 females per group. Data are plotted as violin plots with median and 1st/3rd quartiles and showing all data points. Significantly different groups are indicated by different letters (one-way ANOVA, Tukey’s multiple comparisons test, p<0.05). (**D, G**) Photographs of ovaries (left) and spermathecae with filled sperm (right) from wild type and *Δdum* females 6 days (D) or 12 days (G) post blood-meal with eggs retained. (**E**) Eggs laid by females of the indicated genotype 6 days post-blood-meal. Data are plotted as violin plots with median and 1st/3rd quartiles and showing all data points. Each point represents the eggs laid by a single female (n=47-49/genotype). ns, not significant (Mann-Whitney test). (**F, I, J**) Distribution of eggs laid (F, I) or eggs hatched (J) in wild type and *Δdum* mutant females 6 days (F) and 12 days (I, J) post-blood-meal. Zero values are binned separately for each group. All other bins are groups of 10 starting with [1-10] and with closed/inclusive intervals. (F, I, J) The groups between each genotype for eggs laid and eggs hatched respectively were compared at each of the time points to determine significant difference (Mann-Whitney tests, **** p<0.0001). Distributions in (F) are analyzed from data in (E) and distributions in (I, J) are analyzed from data in (H). (**H**) Number of eggs laid by (top) and hatched from (bottom) single wild type and *Δdum* mutant females 12 days post blood-meal, depicting extended egg retention. Females laying no eggs are depicted by open circles. Lines connect eggs laid by and hatched from the same individual. Larger circles and bold lines represent medians. Boxes show (mean ± S.E.M) of % eggs hatched from each egg retention group, n = 40-45 females/ group.

## Notes

https://doi.org/10.5281/zenodo.5945525

### Competing Interest Statement

The authors have declared no competing interest.

